# Segger: Fast and accurate cell segmentation of imaging-based spatial transcriptomics data

**DOI:** 10.1101/2025.03.14.643160

**Authors:** Elyas Heidari, Andrew Moorman, Dániel Unyi, Nikhita Pasnuri, Gleb Rukhovich, Domenico Calafato, Anna Mathioudaki, Joseph M. Chan, Tal Nawy, Moritz Gerstung, Dana Pe’er, Oliver Stegle

## Abstract

The accurate assignment of transcripts to their cells of origin remains the Achilles heel of imaging-based spatial transcriptomics, despite being critical for nearly all downstream analyses. Current cell segmentation methods are prone to over- and under-segmentation, misassign transcripts to cells, require manual intervention, and suffer from low sensitivity and scalability. We introduce segger, a versatile graph neural network based on a heterogeneous graph representation of individual transcripts and cells, that frames cell segmentation as a transcript-to-cell link prediction task and can leverage single-cell RNA-seq information to improve transcript assignments. On multiple Xenium dataset benchmarks, segger exhibits superior sensitivity and specificity, while requiring orders of magnitude less compute time than existing methods. The user-friendly open-source software implementation has extensive documentation (https://elihei2.github.io/segger_dev/), requires little manual intervention, integrates seamlessly into existing workflows, and enables atlas-scale applications.

## Introduction

Imaging-based spatial transcriptomics (iST) methods offer a powerful means of characterizing single cells within their tissue context. Technologies such as the 10x Genomics Xenium (Janesick et al., 2023), NanoString CosMx (He et al., 2022), and MERFISH (Chen et al., 2015) can profile transcripts from hundreds to thousands of genes at subcellular resolution in a single experiment, often in addition to collecting morphological data. iST data holds great promise for modelling tissues systems, motivating computational innovations for analyzing cell–cell communication and defining cellular niches (Carstens et al., 2024; Dezem et al., 2024; Liu et al., 2024; Moses & Pachter, 2022; Seferbekova et al., 2023). Critically, these modelling advances rely on the cell as a fundamental biological unit, and the basic task of identifying cells and assigning which transcripts belong to them faces serious challenges. While most iST is complemented by nuclear morphology with DAPI, membrane staining is typically lacking. Moreover, the absence of universal cell membrane markers, cellular overlap within the tissue section (Tiesmeyer et al., 2025), transcript diffusion (Ergen & Yosef, 2025) and ambiguity in assigning membrane-localized transcripts all frustrate cell segmentation and transcript assignment even under optimal membrane staining (**Fig. 1a,b**). As a critical first step in iST data analysis, segmentation error can profoundly affect downstream analyses such as cell typing and cell–cell communication inference (Marco Salas et al., 2025; Mitchel et al., 2025; Vasconcelos et al., 2024).

**Figure 1.**
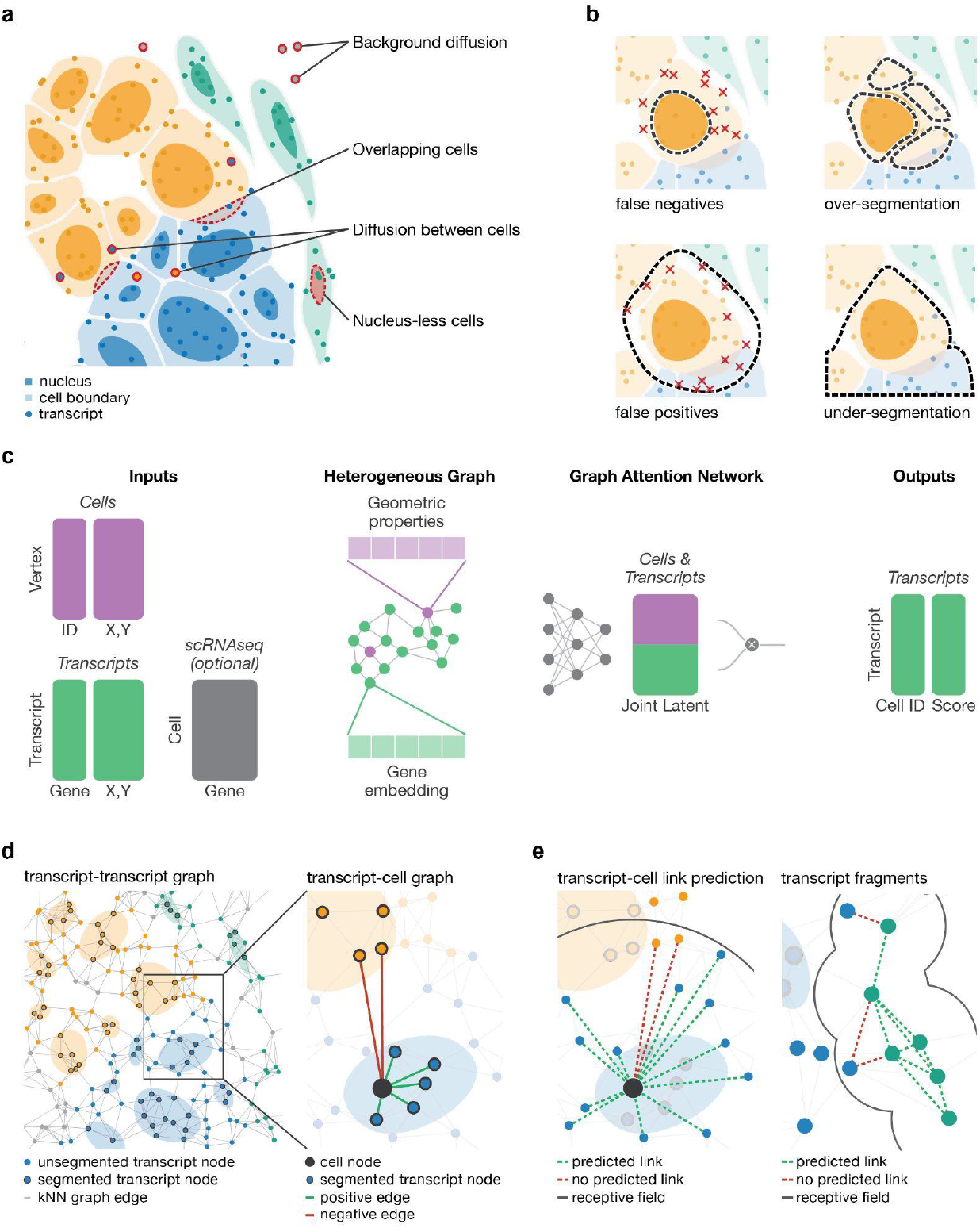
Segger is designed to overcome cell segmentation challenges. **(a)** Accurate cell segmentation is hampered by technical and conceptual challenges posed by iST data, including ambiguity in assigning transcripts to cells due to overlap between cells, transcript diffusion between cells, background diffusion and cells that lack a detectable nucleus. **(b)** Pitfalls and limitations of cell segmentation: Conservative transcript-to-cell assignment may reduce the sensitivity for capturing cytoplasmic transcripts, whereas false positive transcript assignments contaminate the transcriptomic profile of a cell. Segmentation needs to balance over- and under segmentation to avoid splitting biological cells into multiple objects or erroneously combining multiple cells. **(c)** Segger workflow. Segger encodes iST data as a heterogeneous graph of cell and transcript nodes. Cells are represented by geometric features of the staining-derived segmentation masks; genes are represented by a learned or scRNA-seq-informed embedding. An attention-based GNN is trained on initial cell-to-transcript assignments to predict and refine transcript–cell links. **(d)** Heterogeneous graph construction. Initial training instances are generated based on transcript overlap or lack of overlap with cell nucleus boundaries. **(e)** Once trained, segger can be used to refine transcript-to-cell assignments (left), and to cluster unassigned transcripts into ‘fragments’ (right).

Computational approaches have made some progress in segmenting iST data. Classical algorithms such as watershed (Vincent & Soille, 1991) and Voronoi tessellation (Jones et al., 2005) rely solely on transcript density to estimate cell boundaries, making them computationally efficient but inaccurate. Probabilistic approaches such as Baysor and Proseg (Petukhov et al., 2022; Sawada & Honda, 2009) improve on accuracy by learning transcript colocalization patterns, but their computational costs limit practical applicability to larger datasets that are typical with the current technologies. Image segmentation methods such as Cellpose (Riendeau et al., 2024) rapidly segment images based on fluorescent nuclear and membrane marker staining, but cannot assign transcripts to cells; moreover, nuclear segmentation with no or little expansion comes at the cost of lower sensitivity for capturing cytoplasmic transcripts in particular, and most iST data lacks membrane staining. BIDCell (Fu et al., 2024) extends Cellpose by incorporating several custom loss functions based on transcriptomic and morphological data, but there is no clear metric to guide manual tuning of these competing functions. Existing graph-based methods (Defard et al., 2024; Jin et al., 2023) employ graph-based community detection and graph neural networks to group transcripts, but do not leverage the information contained in staining information in a holistic manner. Thus, despite these advances, transcript misassignment remains a major issue (particularly near cell peripheries) (Ergen & Yosef, 2025; Ma et al., 2024; Mitchel et al., 2025; Vasconcelos et al., 2024), and methods that improve accuracy struggle with scalability, underscoring the need for a solution that integrates both spatial and transcriptomic information as well as balancing both speed and accuracy.

To address these challenges, we developed segger, a comprehensive and flexible heterogeneous graph neural network (GNN)–based method (Zhou et al., 2018) that integrates transcript colocalization with nuclear or cell membrane staining data to achieve fast and accurate, 3D-aware segmentation. Segger is an end-to-end segmentation framework that can be applied to any iST data with subcellular resolution. The model only requires a high-confidence initial segmentation, which can be derived from membrane or DAPI nuclear staining captured by all major platforms. The algorithm can either learn a transcript embedding *de novo*, or take on existing embeddings as input, such as those derived from an scRNA-seq reference. We show that segger outperforms leading algorithms on a variety of segmentation tasks.

## Results

Segger frames cell segmentation as a link-prediction task, assigning individual transcripts to cells (**Fig. 1c,d** and **Methods**). As input, it takes iST data, comprising transcripts, represented as points with respective coordinates and transcript identity, and initial cell boundaries, which are either based on nuclear DAPI or cell membrane staining. Optionally, segger can incorporate matching scRNA-seq to inform on gene co-expression patterns. Segger encodes these data as a heterogeneous graph, with transcripts and cells being represented as distinct node types. Transcript–transcript edges reflect the spatial proximity between transcripts and transcript–cell edges encode the assignment of transcripts to cells. During training, segger learns how to correctly associate transcripts to cells using link prediction.

The model is trained based on the observed spatial overlap of transcripts and the staining-derived cell boundaries (positive edges), and negative edges based on transcripts overlap with other neighboring boundaries. Implemented as attention-based GNN (Brody et al., 2021), segger learns a joint latent space in which the proximity of transcript and cell nodes reflects their likelihood of association, capturing gene-based transcript similarity, spatial context, and observed boundary overlap (**Extended Data Fig. 1.1**). Once trained, the model can be used to estimate the evidence for transcript–cell associations (**Fig. 1e**). Segger’s output is a cell segmentation based on the transcripts connected to each cell. In a second grouping step, segger can also assign the remaining unassigned transcripts (which may be due to undetected cells, such as a missing nucleus). In this step, unassigned transcripts are clustered into *fragments* based on their spatial and molecular coherence (**Fig. 1e**). Segger explicitly models these fragments as distinct entities rather than cells.

The open-source segger software is highly optimized for efficient, large-scale segmentation. It employs adaptive tiling (**Extended Data Fig. 1.1**) and efficient task scheduling, supports multi-GPU processing and multi-threading, and comes with data-driven heuristics to reduce the need for manual tuning of parameters and prior knowledge. This ensures that segger delivers both accuracy and speed without compromising flexibility. The software package also includes extensive diagnostic tools, documentation and tutorials (**Methods**).

### Segger enables accurate and efficient segmentation of breast cancer Xenium data

We applied segger and leading alternative cell-segmentation methods to a widely used 10x Genomics Xenium dataset derived from FFPE human breast cancer tissue (313 gene panel) (Janesick et al., 2023) (**Fig. 2, Extended Data Fig 2.1**). Segger was applied using transcript embeddings derived from a breast cancer scRNA-seq atlas (Wu et al., 2021) (**Methods**). For comparison, we also evaluated Baysor (Petukhov et al., 2022), BIDCell (Fu et al., 2024), and the 10X Genomics Xenium software, either with nucleus expansion (hereafter, ‘10x Cell’) or without expansion (‘10x Nucleus’) -methods and analysis strategies that are widely used (Ergen & Yosef, 2025; Hartman & Satija, 2024; Marco Salas et al., 2025; Mitchel et al., 2025). Cell type labels from the scRNA-seq reference were transferred to the segmented cells (**Methods**).

**Figure 2.**
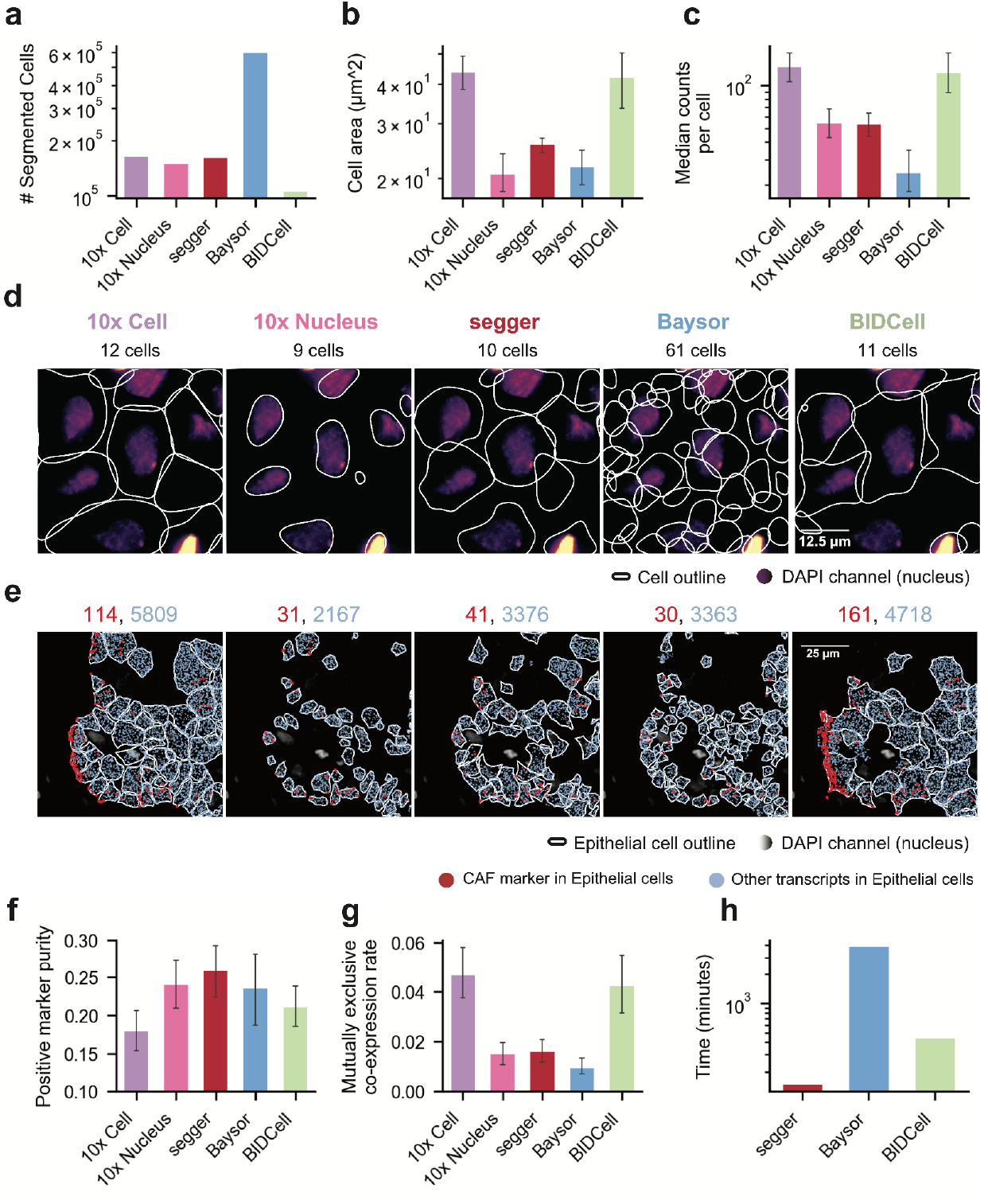
Comparative performance of segmentation methods on the 10x Xenium breast cancer dataset. **(a)** Total number of cells segmented by segger and alternative segmentation tools. Excessive cell counts, as provided by Baysor, are indicative of over segmentation. Cell count from 10x Cell corresponds to the number of stained nuclei. Cells with less than 5 counts have been discarded. **(b)** Median cell area (µm^2^), with segger identifying cell sizes between 10x Cell, and 10x Nucleus. **(c)** Median transcript counts per cell, reflecting the balance between transcript coverage and segmentation. **(d)** Segmentation results in a representative field of view, showing nucleus staining (DAPI channel) alongside transcripts assigned by various segmentation methods. The total count of segmented cells is displayed above each panel. **(e)** Segmentation results in a second representative field of view, analogous to **d** focused on Epithelial cells. Shown are the number of non-CAF transcripts assigned to Epithelial cells (blue) and CAF transcripts (*MMP2, POiSTN, LUM*) falsely assigned to Epithelial cells (red). The total number of transcripts of each category are displayed above each panel. **(f)** Positive marker purity across cells, corresponding to the proportion of correct transcript assignments, as defined from an scRNA-seq atlas. Segger maintains high purity levels, comparable to conservative nucleus-only methods (10X Nucleus). **(g)** Mutually exclusive co-expression rate (MECR) for pairs of exclusive transcripts (N=236 gene pairs, <1% codetection rate in parallel scRNA-seq). Shown is, for each pair of genes, the fraction of cells with false-positive co-expression versus cells with exclusive expression. Baysor, segger, and 10x Nucleus achieve comparable MECRs. **(h)** Empirical compute times evaluated on the whole section for Baysor (without tiling and parallelization), BIDCell, and segger (evaluated on 4 Intel Xeon E5-2620 v4 CPU cores and 4 NVIDIA A100 GPUs). Segger achieves a 30× speedup compared to Baysor and more than a 3 × speedup compared to BIDCell. Metrics in **a-c,f–h** were computed for the full slide; error bars represent interquartile range (IQR).

Segger, 10x Cell and 10x Nucleus detected a comparable number of nucleated cells, with segger yielding cell areas and transcript counts between 10x Cell and 10x Nucleus segmentations (**Fig. 2a–c**). Baysor appeared to over-segment, as it detected many more cells, with few transcripts per cell; indeed, it called up to five times more cells than stained nuclei (**Fig. 2a–d** and **Extended Data Fig. 2.2**). In contrast, BIDCell consistently underestimated the number of cells compared to stained nuclei, resulting in exaggerated cell sizes and transcript counts per cell (**Fig. 2a–d**). Segger’s spatial distribution of cell types is consistent with prior observations (Janesick et al., 2023), and the algorithm preserved expected differences between transcript counts in cancer-associated fibroblasts (CAFs), epithelial cells, immune cells and other cell types (**Extended Data Fig. 2.3**). While segger yields smaller cells than the 10x Cell segmentation on average, a considerable fraction of 10x-assigned transcripts lie far from segmented nuclei and may be wrongly attributed. Indeed, we found that 10x Cell and BIDCell misassigned many more transcripts than segger, particularly at fibroblast–epithelial boundaries, in the representative FOV (**Fig. 2e**).

To investigate transcript misassignment globally, we assessed the positive cell type marker purity (PMP), which provides a measure of the true positive rate based on cell type markers identified from scRNA-seq (**Fig. 2f**), as well as the mutually exclusive co-expression rate (MECR, **Fig 2.g**) as a measure of the false positive rate for each method. Notably, segger achieved the highest PMP across the segmentation methods in this study and a MECR comparable to 10x Nucleus, while maintaining less than half the rate observed in 10x Cell and BIDCell (**Fig. 2g** and **Extended Data Fig. 2.3**).

Thus, whereas Baysor closely matches segger in marker purity and yields the lowest false-positive rate, it frequently over-segments cells (**Fig 2d** and **Extended Data Fig. 2.2**). At the same time BIDCell and 10x Cell produce large, under-segmented cells, generating segmentations with substantial contamination and low marker purity. Segger successfully recovered cytoplasmic transcripts that are missed by 10x Nucleus without sacrificing purity (**Fig. 2e,f**). Segger further provides a seamless segmentation across the entire slide avoiding artifacts observed by methods such as Baysor, which require tiling of large data sets (**Extended Data Fig. 2.1**). The transcriptomic profiles of cells identified by segger also enabled better segregation along expected cell type axes (**Extended Data Fig. 2.4**).

In addition to segmenting cells marked by nucleus staining, segger can be used to explore transcripts that cannot be confidently assigned to any nucleated cell, which in this dataset make up a substantial fraction of transcripts (**Extended Data Fig. 2.5a**). Segger regrouped ∼8.4M of ∼12.2M unassigned transcripts into ∼120K *fragments*. These *fragments* were, on average, smaller than nucleated cells (median 20 transcripts per fragment vs 50 transcripts per cell, **Extended Data Fig. 2.5b,c**), but tended to retain cell-type specific marker expression (**Extended Data Fig. 2.5d**). Given the differences to nucleated cells, we note that the biological relevance of fragments remains to be clarified.

Finally, we evaluated the impact of incorporating gene embeddings on segger’s segmentation performance, finding a clear benefit of incorporating an external scRNA-seq derived reference (**Extended Data Fig. 2.6**). However, even without external information, segger performed competitively against the existing methods (**Extended Data Fig. 2.6**). Additionally, segger’s intrinsically scalable training approach, which utilizes adaptive tiling, ensures compatibility with high-throughput spatial transcriptomics workflows. The resulting speed advantage (**Fig. 2h**) enables the segmentation of large datasets within minutes. Collectively, these results demonstrate how segger is able to achieve high transcript yield and accuracy, while being also practically applicable.

### Validation of segger segmentation using membrane staining

As a further test, we assessed the segmentation of lung adenocarcinoma tissue profiled with the Xenium platform, using an optimized protocol to stain a protein (NaK ATPase) that specifically marks the plasma membrane of epithelial cells (**Methods**). The accurate segmentation of lung cancer tissue is an important but challenging task. Cancer epithelial cells assume a range of novel phenotypes that vary within and between patients and are not represented in healthy reference tissue databases such as the Human Cell Atlas (Laughney et al., 2020). Moreover, lung tumors can exhibit high cellularity with many epithelial–epithelial cell boundaries, whose precise delineation is critical for distinguishing related cell states, in addition to capturing proximal stromal cells. To provide a high-quality segmentation for comparison, we used Cellpose (Riendeau et al., 2024), a method designed to leverage cell-membrane staining. Manual inspection confirmed the suitability of using the Cellpose segmentation as a reference (**Fig. 3a,b**). Segger, 10x and Baysor generated distinct segmentation patterns that strongly impact the accuracy of epithelial cell boundary definitions and transcript assignments (**Fig. 3**). The 10x Cell segmentation tended to over-expand cell boundaries, resulting in larger cell sizes (average ∼180 μm^2^ in 10x; 160 µm^2^ in Cellpose) that were prone to contamination (∼10%) with non-epithelial transcripts (**Fig. 3a,c,e,f** and **Extended Data Fig. 3.1**). Differential expression analysis (FDR < 0.05; DESeq2 (Love 2014)) revealed that only markers of non-epithelial cell types were significantly overrepresented in 10x-segmented epithelial cells compared to Cellpose (**Fig. 3c**). On the other hand, Baysor segmented much more aggressively, more than doubling the number of expected epithelial cells, to give at least two overlapping smaller cells (∼90 μm^2^) per Cellpose epithelial cell on average (**Fig. 3b,f,g** and **Extended Data Fig. 3.2**). This fragmentation contributed to sparse transcript coverage, reduced molecular counts per cell, and distorted uniform manifold approximation and projection (UMAP)-based cell population separation (**Fig. 3h** and **Extended Data Fig. 3.3**). In contrast, segger produced epithelial cell boundaries consistent with the reference, with similar average cell counts to Cellpose ground truth. Segger-segmented cells are marginally smaller than Cellpose (∼140 μm^2^), reflecting a conservative bias against contamination, which can be more problematic for analysis than a slightly lower recall. Indeed, by balancing precision and recall, segger avoids over-segmentation and achieves high transcript assignment rates, while minimizing contamination (∼5%) to accurately demarcate epithelial structures (**Fig. 3e–h**). The only transcript significantly overrepresented in segger compared to Cellpose-segmented cells is an epithelial marker (**Fig. 3d**).

**Figure 3.**
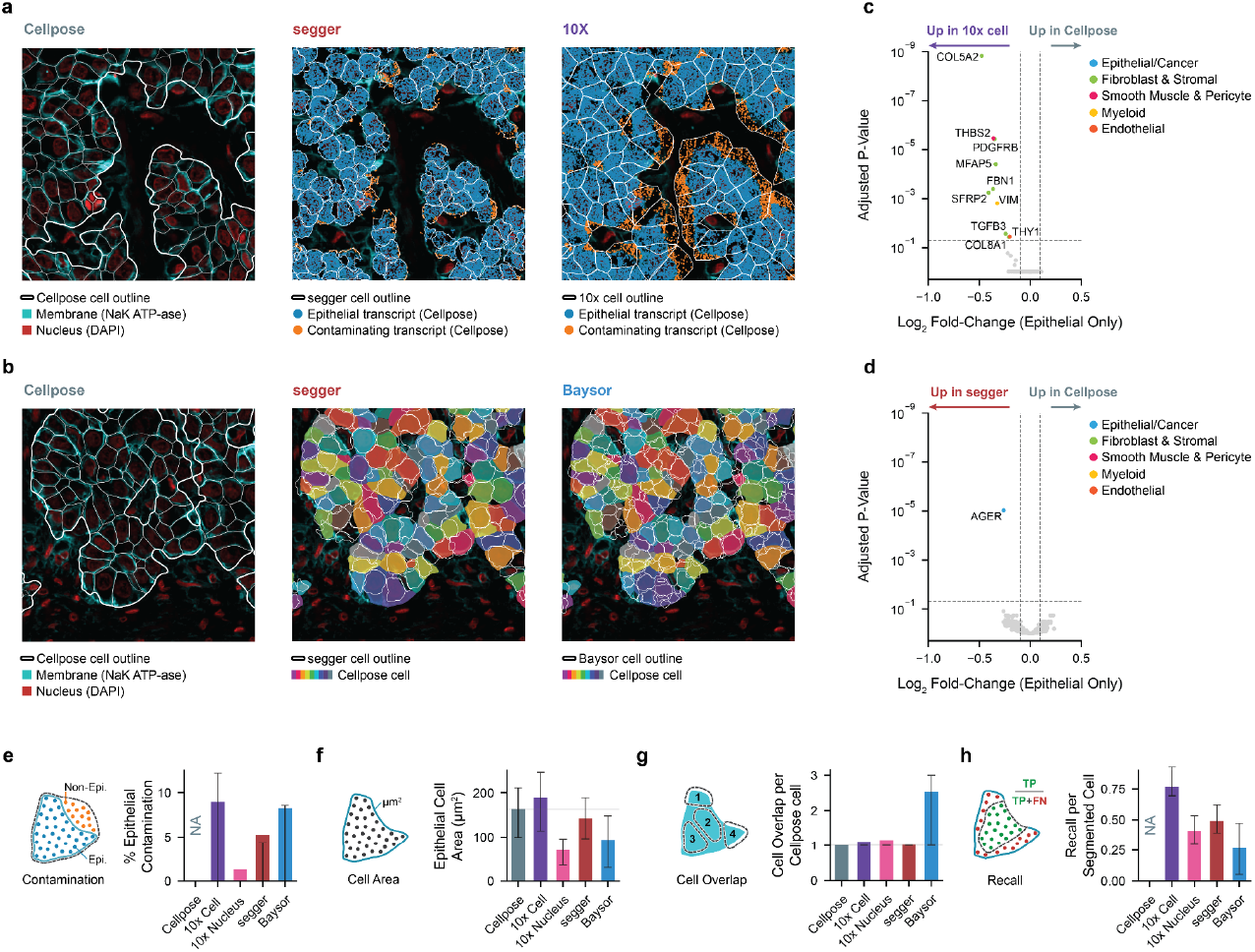
Benchmarking of epithelial cell segmentation in membrane-stained Xenium lung cancer data. **(a)** Cellpose reference segmentation of lung adenocarcinoma epithelial cells, based on cell-membrane stain (left) is used to classify transcripts as epithelial or non-epithelial. Non-epithelial contaminating transcripts (red) are shown in segger (center) and 10x (right) segmentations derived without cell membrane information. **(b)** Cellpose-segmented cells (left), overlaid with segger (center) or Baysor (right) segmented cell boundaries. **(c)** Differential expression analysis to identify transcripts that are differentially captured in 10x versus Cellpose-segmented epithelial cells. Genes annotated by cell type, consistently highlight non-epithelial cell type markers. Shown are adjusted P-values (wald test, Benjamini-Hochberg adjusted) with the horizontal line corresponding to FDR < 0.05. Vertical lines show abs(LogFC) > 0.1 **(d)** Analogous analysis as in **c**, comparing segger segmented cells to Cellpose. The only differentially expressed gene is an epithelial marker (Methods). **(e)** Epithelial transcript contamination, quantified as the fraction of transcripts assigned to epithelial cells by a given method that were not assigned by Cellpose. **(f)** Epithelial transcript recall, quantified as the fraction of Cellpose-assigned epithelial transcripts that were assigned to an overlapping cell by an alternative method, using greedy matching and largest cell overlap. (**g)** Mean epithelial cell area. **(h)** Average number of unique segmented cells that overlap with a Cellpose-segmented epithelial cell. Metrics in **c,d,e–h** were computed for the full slide; error bars represent IQR.

### Segger recovers key cell types and inflammatory niches in human colon data

Given the variation in contamination and recall between methods, we sought to determine the effect of different segmentations on downstream analysis and biological insights, using a healthy human colon Xenium dataset generated by 10x Genomics (10x Genomics 2024). UMAP projection of count matrices derived from segger segmentation of these data revealed the expected cell types, along with a distinct neutrophil-like subpopulation that robustly expresses immune-associated inflammatory response markers such as *IL1B* (**Fig. 4a–c** and **Extended Data Fig 4.1a**). These observations are supported by histology showing neutrophil accumulation within the colonic lumen (**Extended Data Fig. 4.1b**)—a known phenomenon during acute or localized colonic injury (Azcutia et al., 2022).

**Figure 4.**
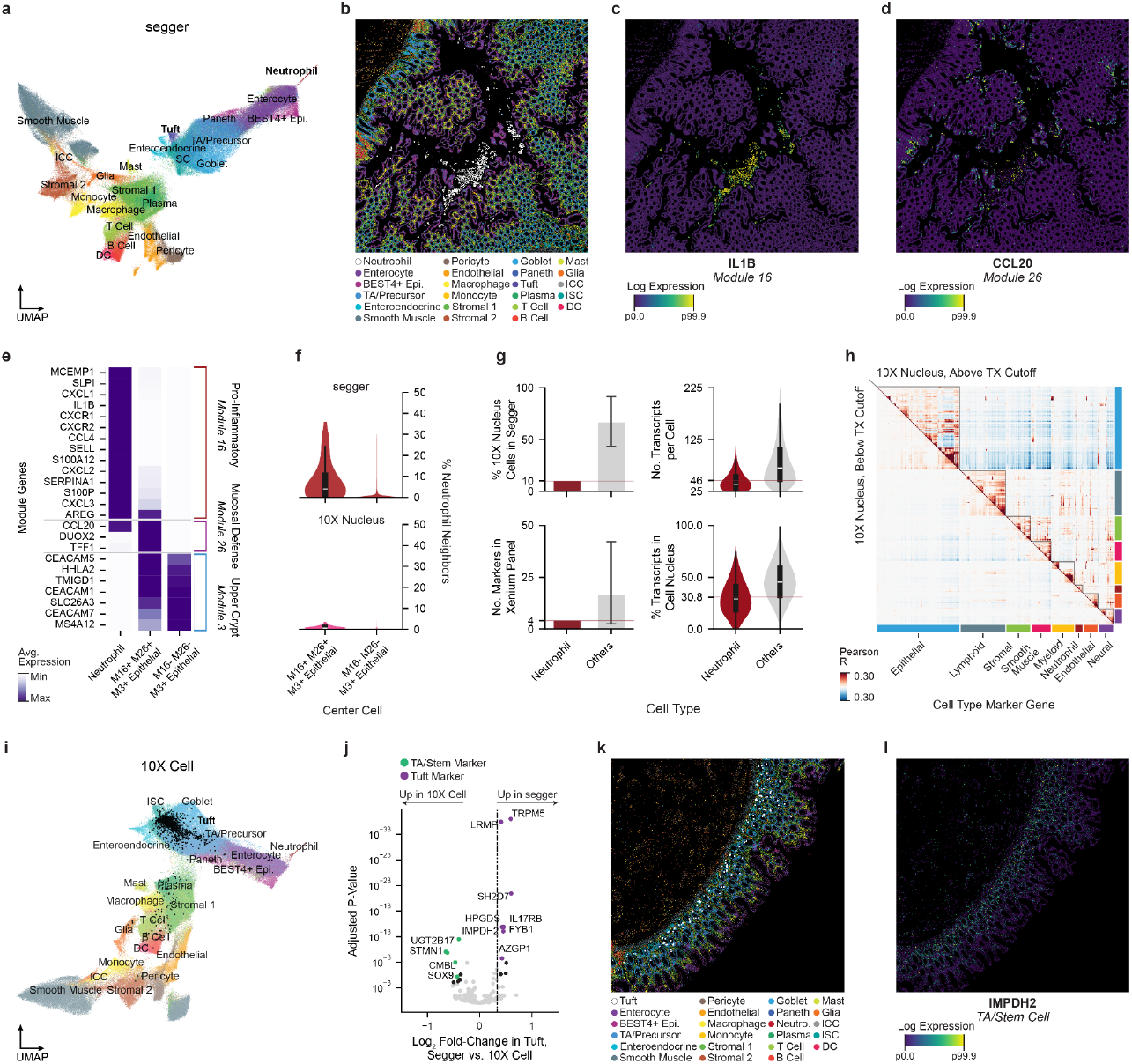
Segger recovers cytoplasmic cell types and preserves neighborhood structure in normal human colon. **(a)** UMAP projection of segger-segmented cells, displaying distinct cell type populations, including neutrophils, tuft cells, enterocytes, and stromal populations. **(b)** Localization of cell types in a representative field of view (Methods). Epi., epithelial; TA, transit amplifying stem cellI; ICC, interstitial cell of Cajal; ISC, intestinal stem cell; DC, dendritic cell. (**c**) Normalized log-scale expression of neutrophil marker *IL1B*, belonging to Hotspot module 16. **(d)** Normalized log-scale expression of immune cell recruitment chemokine *CCL20*, belonging to Hotspot module 26. **(e)** Relative average expression of genes from Hotspot modules 16, 26, and 3 across three groups: neutrophils, neighboring epithelial cells enriched for modules 16, 26, and 3, and epithelial cells of the upper crypt expressing only Module 3. **(f)** Relative recovery of cell states in segger and 10x Nucleus segmentation, highlighting differences in detection rates. **(g)** Lower neutrophil cell recall by 10x Nucleus (upper left) reflects the low total RNA content (upper right), low nuclear transcript proportion (bottom right) and few panel markers (bottom left) for this cell type. **(h)** Gene–gene correlation matrices of all 10x Nucleus-segmented cells above (upper right) and below (lower left) a 25-transcript-per-cell cutoff, showing that lowering the threshold in an attempt to improve sensitivity would only add data with little to no cell-type signal; it would not recover missed cell types. **(i)** UMAP projection of 10x Cell-segmented data, showing the clustering of different cell types. **(j)** Differential gene expression analysis comparing segger and 10x Cell-segmented tuft cells, highlighting cell-type-specific markers. **(k)** Tuft and TA/stem cells share overlapping spatial domains in colon tissue. **(l)** Spatial distribution of TA/stem cell marker IMPDH2.

To explore the regions surrounding neutrophil infiltration, we applied Hotspot (DeTomaso & Yosef, 2021) to identify modules of genes with spatially correlated expression (**Extended Data Fig. 4.1c, Supplementary Table 4** and **Methods**). Of these, genes from Module 16 are upregulated in neutrophils, whereas the most correlated module (Module 26) is primarily expressed in epithelial cells and contains mucosal defense genes such as *CCL20*, a chemokine involved in the recruitment and migration of immune cells including neutrophils (Danne et al., 2023) (**Fig. 4c–e**). Module 26+ cells form a distinct subpopulation within the upper crypt, which is itself characterized by Module 3 gene expression (**Fig. 4e** and **Extended Data Fig. 4.1d,e**). An analysis of cell type proportions within cell neighborhoods (*k* = 200 nearest neighbors; **Methods**) revealed that neutrophils make up a higher proportion of the surrounding cells for Module 16/26+ epithelial cells compared to Module 16/26-cells, suggesting an injury-response “axis” between inflammatory neutrophils and these cells.

Nuclear segmentation is a common choice for iST data, as DAPI staining is robust and the corresponding transcript assignments are considered faithful and free of contamination. However, the loss of cytoplasmic signal, which contains the majority of cellular transcripts (typically on the order of 5-fold more than the nucleus (Abdelmoez et al., 2018)), can reduce sensitivity to the point of missing important biology. In the colon data, the same analysis using 10x Nucleus segmentation failed to identify variation in neutrophil abundance between Module 16/26-positive and negative cells (**Fig. 4f**). Further investigation revealed that neutrophils are severely underrepresented in this segmentation owing to their small size and predominantly cytoplasmic transcript localization, in addition to having few markers in the assay panel (**Fig. 4g**). The resulting low transcript capture (recall) per neutrophil makes these cells difficult to distinguish from noise (**Fig. 4h** and **Extended Data Fig 4.1f**). By contrast, segger captures enough cytoplasmic transcripts to robustly identify neutrophils, without compromising gene covariance structure, a key indicator of biological coherence in data (**Extended Data Fig. 4.1g**).

We next compared segger to 10x Cell, which has better recall but is also subject to higher levels of transcript contamination. We found differences particularly in epithelial populations, such as tuft cells, a specialized lineage involved in chemosensation and immune signaling. While segger accurately captures tuft cells and their boundaries, 10x Cell frequently misassigns these cells, blending their transcriptional profiles with adjacent stem-like populations (**Fig. 4i**). The 10x Cell segmentation contaminates tuft cell profiles with stem cell markers such as *LGR5* and *SOX9*, whereas segger maintains the integrity of tuft markers such as *TRPM5* and *POU2F3* (**Fig. 4j,** FDR < 1e-3; DESeq2 (Love 2014)). Spatial overlap between tuft and stem or transit-amplifying marker (*IMPDH2*) localization explains the origin of this discrepancy in the 10x Cell segmentation, which is prone to over-expanding cell boundaries (**Fig. 4k,l**).

Our comparisons demonstrate that segger achieves an effective balance between assigning sufficient cytoplasmic transcripts to capture biological entities with low signal, such as neutrophils and their potential niche interactions, while avoiding transcript contamination that can greatly mislead cell-state analysis.

### Discussion

The segger GNN approach uniquely frames cell segmentation as a supervised link prediction task, significantly enhancing segmentation accuracy and speed for spatial transcriptomics data. Across three Xenium datasets—including a lung adenocarcinoma dataset with membrane staining as ground-truth cell boundaries—segger achieved well-balanced segmentation, effectively extending cells beyond nuclei while minimizing contamination and fragmentation. In contrast, we demonstrated that existing tools tend to over-segment, resulting in inflated cell counts and poor transcript recall; restrict segmentation to nuclei, causing transcript sparsity and reduced sensitivity; or under-segment, producing expanded cell boundaries that mix transcripts from multiple cells.

Poor segmentation has significant implications for downstream biological analysis. In human colon data, we observed that nuclear segmentation without expansion, which fails to assign cytoplasmic transcripts, completely eliminates neutrophil detection and prevents characterization of inflammatory microenvironment responses. Segger overcomes these limitations by recovering cytoplasmic transcripts, but critically—unlike default 10x Cell segmentation—this improved sensitivity does not come at the cost of increased contamination. While some residual contamination is inevitable regardless of the segmentation method, recent studies suggest that these signals originate from spatially overlapping cells, partial cell fragments within the imaging plane, and molecular diffusion during tissue preparation (Ergen & Yosef, 2025; Hartman & Satija, 2024; Mitchel et al., 2025; Tiesmeyer et al., 2025).

Segger is built on a flexible modeling framework, creating opportunities for future extensions. (Ergen & Yosef, 2025). In principle, segger’s gene embeddings already provide a foundation for incorporating prior information from scRNA-seq for this task. Currently, segger does not explicitly filter out contamination from transcripts overlapping straining-derived nucleus boundaries, as these edges are treated as positive connections during model training. Enhancing segger’s ability to address this source of contamination would not only refine segmentation accuracy but also improve the identification of biologically meaningful cell fragments—spatially and transcriptionally coherent transcript groups that cannot be confidently assigned to a single cell boundary.

Once trained, the segger model can be readily transferred to new tissue sections and datasets. Future work will explore its application to large collections of iST datasets, allowing segger to learn more robust and generalizable rules for transcript-to-cell assignment. Finally, while segger is already applicable to any single-molecule technology with membrane or nucleus staining information, the model could be further extended to other platforms, such as Visium HD (Nagendran et al., 2023), which aligns well with the graph-based modeling approach employed in segger.

## Methods

Methods and supplementary notes are provided as a separate file.

## Data availability

The breast cancer dataset was obtained from the 10x Genomics website on May 6, 2024: https://www.10xgenomics.com/products/xenium-in-situ/preview-dataset-human-breast. The healthy colon dataset was downloaded from the 10x Genomics website on January 15, 2025:https://www.10xgenomics.com/datasets/ffpe-human-colorectal-cancer-data-with-human-immuno-oncology-profiling-panel-and-custom-add-on-1-standard. Newly generated data, including intermediate outputs, segmentation results for segger, Baysor, BIDCell, and CellPose, as well as the trained models required for reproducing the analyses in this study, are publicly available on our S3 bucket:

- Breast cancer dataset: https://dp-lab-data-public.s3.us-east-1.amazonaws.com/segger/xenium_breast.tar
- NSCLC dataset: https://dp-lab-data-public.s3.us-east-1.amazonaws.com/segger/xenium_nsclc.tar.gz
- Colon dataset: https://dp-lab-data-public.s3.us-east-1.amazonaws.com/segger/xenium_colon.tar.gz

## Code availability

Segger is available as an open source python package at: https://github.com/PMBio/segger. Scripts and custom code for data analysis related to this manuscript are available at https://github.com/dpeerlab/segger-analysis. Online documentation and tutorial for the package are available at: https://elihei2.github.io/segger_dev.

## Author contributions

E.H. conceived the initial segger model. E.H., A.Mo., M.G., D.P. & O.S. conceived the validation strategy and further extensions of segger. E.H. & A.Mo. developed the segger software with contributions from D.U. G.R. implemented the cell mask boundary generation algorithm. E.H. conducted analyses on the Breast cancer dataset with input from G.R. & A.Mo. A.Mo. conducted analyses on the NSCLC and colon datasets with input from E.H. J.C. collected the clinical NSCLC sample. N.P. processed the sample using the Xenium platform, optimized antibody staining, and performed IF membrane staining. E.H., A.Mo., T.N., M.G., D.P., and O.S. wrote the manuscript with contributions from all authors. M.G., D.P., and O.S. supervised the study.

## Acknowledgments

We thank S. Rutz, D. Dimitrov, and L. Marconato for contributions to manuscript writing, and L. Marconato for input on extending segger to use SpatialData. We acknowledge S. Kher and F. Hyel for initiating extensions to the nf-core module. We are grateful to the scverse community and F. Theis for valuable insights on scaling and software engineering. We also thank the organizers and participants of SpaceHack Germany (2022 and 2024) for input on software usability and extensions leveraging the Xenium segmentation kit. We thank T. Dougherty, N. Pasnuri, and A. Jimenez-Sanchez for their contributions to idea development and our understanding of *segger* during the Pe’er Lab retreat hackathon. We also thank R. Niec for her insights on histological evaluation and interpretation of neutrophil accumulation in the colon dataset. This work was supported by the German Bundesministerium für Bildung und Forschung (BMBF) through the project “DeepSC2 - Bioinformatische Methoden auf der Basis von tiefen Netzen für die Analyse einzelner Zellen in der Onkologie” [031L069A] and the European Union project RRF-2.3.1-21-2022-00004 within the framework of the Artificial Intelligence National Laboratory. We also gratefully acknowledge support from the National Institutes of Health (NIH) cancer center support grant P30 CA08748 (D.P.) and U54 CA274492 (D.P), the Gerry Metastasis and Tumor Ecosystems Center, and the Single-cell Analytics Innovation Lab at Memorial Sloan Kettering Cancer Center. D.P. is a Howard Hughes Medical Institute investigator. M.G. acknowledges support through the HEROES-AYA consortium within the National Decade Against Cancer of the German Federal Ministry of Education and Research (01KD2207).

