## Supplementary material for "Segger: Fast and accurate cell segmentation of imaging-based spatial transcriptomics data": Online methods and supplementary notes

### segger: Online Methods and Supplementary Notes

#### Contents

**1 segger model** 4
- 1.1 Overview 4
- 1.2 Heterogeneous graph construction 4
  - 1.2.1 Graph nodes 4
  - 1.2.2 Graph edges 5
  - 1.2.3 Tiling Strategy 5
- 1.3 GNN model and training 5
  - 1.3.1 Model architecture 5
  - 1.3.2 Loss function and training 6
- 1.4 Prediction and segmentation 7
  - 1.4.1 Assigning transcripts to cells 7
  - 1.4.2 Grouping unassigned transcripts into fragments 7
- 1.5 Relationship between segger and existing cell segmentation methods 8

**2 Heuristics and parameter selection guidelines** 8
- 2.1 Tiling parameters 8
- 2.2 GNN model parameters 8
- 2.3 Prediction of transcript–cell links 9
- 2.4 Segger confidence score cut-off for transcript–cell link prediction 9
- 2.5 Transcript–transcript receptive fields for identifying fragments 9

**3 Data analysis** 10
- 3.1 Default 10x cell segmentation 10
- 3.2 Breast cancer Xenium dataset 10
  - 3.2.1 Dataset description 10
  - 3.2.2 segger 10
  - 3.2.3 Baysor 10
  - 3.2.4 BIDCell 10
  - 3.2.5 Preprocessing of segmented cells 11
  - 3.2.6 Cell type marker genes for benchmarking from scRNA-seq 11
  - 3.2.7 Mutually Exclusive Co-expression Rate (MECR) 11
  - 3.2.8 Positive Marker Purity (PMP) 11
  - 3.2.9 Compute time and memory benchmarking 12
- 3.3 NSCLC Xenium dataset 12

**E *segger's* step-by-step workflows** **36**

### 1 segger model

#### 1.1 Overview

Segger is a graph neural network (GNN)-based method that formulates cell segmentation in imaging-based spatial transcriptomics (iST) data as a problem of predicting links between transcripts and cells. The model builds on the intuition that nearby transcripts likely originate from the same cell, by exchanging information locally across a heterogeneous graph: (1) a transcript–transcript graph that connects neighboring transcripts and (2) a transcript–cell graph that reflects an initial high-confidence assignment of a subset of transcripts to putative cells—most commonly, transcripts that overlap nucleus staining. Segger infers a model of plausible transcript–cell associations by propagating information along this heterogeneous graph. Once trained, the model can be used to refine the initial transcript–cell associations, as well as to link previously unassigned transcripts to cells (**Fig. 1**).

The inputs to segger consist of transcripts and cells. Cells are represented by 2D polygon masks, such as those derived from cell nucleus staining. Transcripts are represented by X-Y-Z coordinate positions and gene identifiers. Optionally, the model can leverage scRNA-seq data to inform the assignment of transcripts to cells. The segger worfklow (**Fig. 1c**) comprises three main steps, which are elaborated in the following sections: (1) construction of a heterogeneous graph from the input iST data, (2) training of a GNN based on the heterogeneous graph, and (3) prediction of transcript–cell assignments. Guidelines on how to set model parameters are discussed in Section 2.

#### 1.2 Heterogeneous graph construction

segger models an image-based ST dataset as a heterogeneous graph $\mathcal{G} = (\mathcal{V}, \mathcal{E})$ of two types of nodes and edges. Nodes $\mathcal{V}$ comprise transcripts $\mathcal{T}$ and cells $\mathcal{C}$, and edges $\mathcal{E}$ reflect the spatial colocalization of transcripts, $\mathcal{E}_{TT}$, and an initial assignment of a subset of transcripts to cells, $\mathcal{E}_{TC}$.

##### 1.2.1 Graph nodes

**Transcript nodes** Transcript nodes $t_i \in \mathcal{T}$ consist of measured transcripts which possess a gene label $g_i$ and spatial position $p_i \in \mathbb{R}^3$. Each transcript node is associated with an embedding vector $e_g$ of dimension $d_{in}$ that is indexed by its gene label $g$, such that all transcripts of the same gene share a common representation. If no predefined embeddings are available, these vectors are randomly initialized from $\mathcal{N}(0, 1)$. Alternatively, transcript nodes can be initialized with biologically informed embeddings which capture gene similarities from the outset. This provides segger with a prior on plausible co-expression patterns within the same cell. In this manuscript, for each iST dataset, we construct embedding vectors from relevant annotated single-cell RNA sequencing (scRNA-seq) data, where each vector represents gene expression abundance across cell types. Specifically, each gene $g$ is assigned an embedding vector $e_g$ according to the proportion of cells that exhibit nonzero expression for each cell type $l \in L$ in the scRNA-seq data:

$$e_{g,l} = \frac{\sum_{i \in C_l} \mathbb{1}(x_{i,g} > 0)}{|C_l|} \quad (1)$$

where $x_{i,g}$ represents the scRNA-seq count of $g$ in cell $i$, $\mathbb{1}(x_{i,g} > 0)$ is an indicator function that equals 1 if the gene is expressed in the given cell and 0 otherwise, and $C_l$ denotes the set of cells assigned to cell type $l$.

To generate these embeddings in the segger package, users can provide an scRNA-seq input file via `scrnaseq_file` and specify the cell type annotation column with `celltype_column` . See Section 2 and Appendix C for details.

**Cell nodes** Cell nodes $c_i \in \mathcal{C}$ represent segmented cells, each defined by a boundary polygon $B_i \subset \mathbb{R}^2$. These boundary polygons originate from a user-provided partial segmentation, which assumes transcript assignments are incomplete but predominantly accurate. For many users, we assume this segmentation will be based on DAPI-stained nuclei, as provided by platforms such as Xenium (all cell nodes in this manuscript), MERSCOPE, or CosMx.

Each cell node is assigned a feature vector comprising geometrical properties of its boundary polygon $B_i$, which provide cues on cell similarity. Morphological features have been widely shown to capture both broad (e.g., cell type) and subtle phenotypic differences among cells [1, 2]. Here, we leverage this to refine transcript-to-cell assignment for morphologically distinct cells based on four properties: area $A(B_i)$, convexity $C(B_i)$, elongation $E(B_i)$, and circularity $\Gamma(B_i)$:

$$C(B_i) = \frac{A(\text{Conv}(B_i))}{A(B_i)} \qquad E(B_i) = \frac{A(\text{MBR}(B_i))}{A(\text{Env}(B_i))} \qquad \Gamma(B_i) = \frac{A(B_i)}{r_{\text{min}}(B_i)^2} \quad (2)$$

where Conv is the convex hull, MBR is the minimum bounding rectangle, Env is the envelope (the minimum bounding rectangle with sides parallel to the coordinate axes), and $r_{min}$ is the minimum bounding radius. All properties are computed with the package *geopandas* (v1.0.1).

##### 1.2.2 Graph edges

**Transcript–transcript edges** Transcript–transcript edges $\mathcal{E}_{TT}$ capture spatial colocalization and are constructed using a radius-restricted $k$-nearest-neighbor ($k$ NN) graph. Each node connects to at most $k$ nearest neighbors based on Euclidean distance, with edges $(t_i, t_j)$ included only if $d(p_i, p_j) \leq r$, restricting connections to a set radius, $r$. The neighbor constraint prevents excessive edges between transcripts in dense regions while the distance constraint ensures that connections in sparse areas remain localized and do not extend beyond a reasonable cell boundary. In the segger package, these parameters are represented as `k_tx` and `dist_tx` .

**Transcript–cell edges** Transcript–cell edges associate each transcript with the cell to which it is assigned. For training, we construct an "incomplete" version of this graph from the high-confidence segmentation. This is done by assigning transcripts to cells whose boundaries contains the transcripts' spatial coordinates. Formally, for each pair $(t_i, c_j) \in \mathcal{T} \times \mathcal{C}$, edges are defined as:

$$\mathcal{E}_{TC} = \{(t_i, c_j) \mid (p_{i,x}, p_{i,y}) \in \text{int}(B_j)\}, \tag{3}$$

where $(p_{i,x}, p_{i,y})$ denotes the $(x, y)$ coordinates of transcript $t_i$, and $\text{int}(B_j)$ denotes the interior of the boundary polygon associated with cell $c_j$. Once the *segger* model is trained, we then perform link prediction to infer additional transcript-cell edges which were not present in the initial graph (see section 1.4.1 for more details).

In practice, many iST platforms pre-compute transcript-to-cell assignments and provide them directly as transcript labels. When provided, this removes the need for *segger* to compute these assignments using costly geometric comparisons. In this manuscript, we derive the initial transcript–cell graph from the assignment of transcripts to their corresponding *cell id*, which is provided as a transcript label in Xenium datasets, and restrict edges to those within the nuclear segmentation.

##### 1.2.3 Tiling Strategy

Because training on a complete iST graph is computationally intractable and segmentation is inherently local, we use a tree-based tiling strategy to spatially partition the dataset into smaller tiles, and construct graphs independently for each. Tiles are sized to balance transcript counts (versus cell counts), as transcripts vastly outnumber cells and primarily determine graph size and computational cost. This ensures even workload distribution and prevents bottlenecks during graph construction and training.

Our tiling strategy recursively partitions the dataset along the spatial dimension, $X$ or $Y$, with the largest range of transcript positions, splitting at the median position. This process continues until $N$ regions are reached, with each region's data loaded into memory as a separate job. Regions are then further subdivided using the same approach into smaller spatial tiles, each containing approximately $K$ transcripts. For example, when tiling the Xenium colon dataset (see section 3.4), we choose $K = 50,000$, yielding tiles with dimensions of $223 \times 236 \mu m$ and 317 cells on average. To prevent cells from being split across tiles, adjacent regions are expanded by an overlapping margin set by a user-defined parameter. This margin must be large enough to fully contain any cell that would otherwise be split at the boundary (see Section 2 for recommendations). Only cells entirely within the expanded region are retained in the tile. Finally, tiles are stored as separate sets of training, validation, and testing tiles based on the user's parameter choices for the relative size of each set.

Although we use tiling to accelerate training, the mini batch training scheme of *segger* eventually considers all tiles during training, thus learning a single "global" model that is applied to all tiles during prediction. This model captures patterns across the entire dataset, thereby avoiding boundary effects. In contrast, the tiled application of Baysor generates independent models based solely on the data within each tile, leading to inconsistencies at tile boundaries when models constructed from adjacent tiles differ.

#### 1.3 GNN model and training

Building on the heterogeneous graph, segger uses a supervised GNN to infer transcript-to-cell assignments. By incorporating cell and transcript node features, the model passes evidence on transcript-to-cell assignments following a "guilt by association" approach. For instance, if transcript A is spatially proximal and molecularly similar (close in the gene embedding space) to transcript B—and transcript B is linked to a given cell—this suggests that transcript A also belongs to that cell [3, 4, 5] (see Appendix A and **Extended Data Fig. 1.1b**). The GNN captures richer dependencies than guilt by association, however. The model propagates features associated with individual cells and transcripts, and integrates the evidence from multiple transcripts to predict likely transcript-cell associations.

##### 1.3.1 Model architecture

Node features are processed in hidden layers that comprise graph attention and linear transformations (**Extended Data Fig. 1.1b**). The first layer consist of two parallel fully connected linear layers with output size $d_1$

(parameter `init_emb` ), one for transcript nodes, and one for cell nodes. This is followed by a stack of additional GATv2Conv [6] layers, each with output dimension $d_2$ (set by `hidden_channels` . Output from these layers is aggregated and passed through a Leaky ReLU activation [7]. The concluding layer is a fully connected linear layers that maps both transcript and cell nodes to a common $d_3$-dimensional latent space, which serves as input to the loss function for training (with $d_3$ specified by `out_channels` ).

We use multi-head GATs as the core building blocks of segger because of their ability to adaptively weigh and prioritize informative interactions [4] (Appendix A.2). GATv2Conv further enhances this ability by providing flexible, order-invariant attention, which has been demonstrated to improve feature aggregation across heterogeneous components [8]. In the context of segger, the GATs equip the model with flexibility to balance and correct evidence from the transcript–transcript and transcript–cell graphs. The number of attention heads across all GATv2Conv operators is the same and controlled by `heads` .

Intuitively, through *segger*'s graph neural network, each transcript gathers information from nearby transcripts and from overlapping cell nodes. In practical terms, rather than each transcript or cell making an isolated decision based solely on its own features, every transcript "listens" to its nearby neighbors and to overlapping cell nodes. In particular, the attention mechanism, within GATv2Conv layers, assigns weights to these incoming messages, effectively highlighting the more informative signals—such as a strong gene identity match or clear morphological cues from a cell boundary—while down-weighting less relevant ones. However, this process must be carefully moderated, as excessive message passing can lead to oversmoothing, where distinctive features become diluted. Consequently, *segger* typically uses a moderate number of layers to enhance transcript-to-cell assignment while preserving the unique identity of each node (see Section 2).

##### 1.3.2 Loss function and training

The final $d_3$ dimensional layer in segger is linked to a binary classification loss which, given appropriate positive and negative training instances, differentiates between true and false transcript–cell edges.

**Generation of positive and negative training examples** For training, positive and negative examples are derived locally for each tile (c.f. Section 1.2.3). The set of positives edges, $\mathcal{E}_+$, are the same as the edges from the prior segmentation graph $\mathcal{E}_{TC}$. Negative edges, $\mathcal{E}_-$, are obtained from transcripts with evidence for assignment to a non-matching cell. For example, under nuclear segmentation, a negative edge represents a nuclear transcript of one cell paired with the nucleus of a different cell. Formally, the set of negative edges is defined as:

$$\mathcal{E}_- = \{(t_i, c_j) \in \mathcal{T}_C \times \mathcal{C} \mid (t_i, c_j) \notin \mathcal{E}_+\}, \tag{4}$$

where $\mathcal{T}_C \subset \mathcal{T}$ is the set of transcripts that have at least one positive edge.

Since $|E_-|$ by far exceeds $|E_+|$, a random subset of $E_-$ with size proportional to $|E_+|$ is drawn, thereby controlling class imbalance and reducing computational cost. Notably, since negative edges are sampled locally within each tile, this approach naturally biases the selection toward harder negatives—cells and transcripts that are proximal and therefore more likely to share molecular characteristics.

**Binary classification loss** The segger GNN assigns each transcript $t_i$ and cell $c_j$ a representation in the final layer with $d_3$ dimensions, denoted $f(t_i)$ and $f(c_j)$ respectively. A sigmoid loss function is employed to connect the inner product of these feature representations, $s_{ij} = \langle f(t_i), f(c_j)\rangle$, to the likelihood of a positive edge between these nodes [9].

$$\hat{y}_{ij} = \sigma(s_{ij}) = \frac{1}{1 + e^{-s_{ij}}}. \tag{5}$$

These probability values also serve as the segger confidence score (ranging from 0 to 1), which are used for downstream prediction.

Model training is implemented using a supervised classification objective based on the transcript–cell labeled edges $E_- \cup E_+$ with training labels $y_{ij} \in \{0, 1\}$, as defined above. The binary cross-entropy loss is defined as

$$\mathcal{L} = - \sum_{(t_i, c_j) \in E_{tc}} \left[ y_{ij} \log \sigma(s_{ij}) + (1 - y_{ij}) \log\bigl(1 - \sigma(s_{ij})\bigr) \right].$$

**Mini-batch training and monitoring of convergence** Segger employs a distributed mini-batch scheme to train the GNN model [10]. After constructing and splitting tiles into training, validation, and test sets, the training and validation tiles are separately grouped, uniformly and by chance, into mini-batches. These mini-batches are then treated as a single graph, by simply combining their sets of nodes and edges, and fed into the model. The model is trained over multiple epochs, with link prediction performance monitored on the validation tiles using the Area Under the Receiver Operating Characteristic (AUROC) and the F1 score.

#### 1.4 Prediction and segmentation

After training, segger gives rise to a common latent space between transcript and cell in which the inner-product of the embeddings yields a predictive score over transcript–to-cell associations.

##### 1.4.1 Assigning transcripts to cells

Cell segmentation is achieved by assigning each transcript to the cell with the highest similarity score computed from the final node embeddings. Rather than considering all cells globally, each transcript $t_i$evaluates only a subset of candidate cells within its localized *receptive field*. To this end, segger constructs a receptive-field transcript-to-cell graph, $\mathcal{G}_{\text{t-c}}$, based on a radius-restricted $k$ NN search. This search is parameterized by $K_c$ ( `k_bd` in the software), which limits the number of nearest boundary-associated cells per transcript, and by $r_c$ ( `dist_bd` ), which defines the maximum Euclidean distance for including a candidate cell. Formally, the set of candidate cells for transcript $t_i$ is given by

$$\mathcal{C}(t_i) = \{c_j \mid d(t_i, c_j) \leq r_c,\ |\mathcal{C}(t_i)| \leq K_c\}\,.$$

For each candidate cell $c_j \in \mathcal{C}(t_i)$ with embedding $f'(c_j) \in \mathbb{R}^{d_3}$ and transcript $t_i$ with embedding $f(t_i) \in \mathbb{R}^{d_3}$, the similarity score is computed as

$$s_{ij} = \langle f(t_i), f'(c_j)\rangle.$$

The maximum similarity score for transcript $t_i$ is then determined by

$$s_i^{\max} = \max_{c_j \in \mathcal{C}(t_i)} s_{ij},$$

with the corresponding cell assignment given by

$$c_{j^*} = \arg \max_{c_j \in \mathcal{C}(t_i)} s_{ij}.$$

A transcript is assigned to cell $c_{j^*}$ only if $s_i^{\max} > \theta$, where $\theta$ is a user-defined confidence threshold set by `score_cut` . This threshold ensures that only assignments with sufficient confidence are accepted. Transcripts failing to meet this criterion remain unassigned and are later grouped into *fragments* based on their spatial and molecular coherence, thereby capturing critical cytoplasmic information and preserving the overall biological context.

##### 1.4.2 Grouping unassigned transcripts into fragments

Transcripts with a maximum similarity score below the threshold ($s_i^{\max} < \theta$) or without any nearby candidate cell are labeled as *unassigned*. To recover additional information, segger clusters these unassigned transcripts into transcriptionally coherent *fragments* based on spatial and molecular similarity.

This is achieved by constructing a receptive-field transcript-to-transcript graph, $\mathcal{G}_{\text{t-t}}$, using a radius-restricted $k$ NN (fe) search on the unassigned transcripts. Two parameters control this process: $K_t$ ( `k_tx` ), which limits the number of nearest neighbors per transcript, and $r_t$ ( `dist_tx` ), which defines the maximum Euclidean distance for a neighbor.

Let

$$U = \{t_i \mid s_i^{\max} < \theta \text{ or } \mathcal{R}(t_i) = \varnothing\}$$

denote the set of unassigned transcripts. For each $t_i \in U$, its neighborhood is defined as

$$\mathcal{T}(t_i) = \{t_k \in U \mid d(t_i, t_k) \leq r_t\} \quad \text{with } |\mathcal{T}(t_i)| \leq K_t.$$

Pairwise similarity scores are computed via the inner product:

$$s_{ij}^{\text{t}} = \langle f(t_i), f(t_j)\rangle.$$

An edge $(t_i, t_j)$ is added to the graph $G_U = (U, E_U)$ if $t_j \in \mathcal{T}(t_i)$ and $s_{ij}^{\text{t}} \geq \lambda$.

Finally, $G_U$ is partitioned into connected components using standard graph partitioning techniques [11], with each component representing a fragment—a cluster of unassigned transcripts that likely correspond to cell fragments or partial cell profiles.

Fragments can recover a potentially large set of transcripts that are missed by methods solely relying on nuclear or membrane markers [12]. This approach minimizes over-segmentation, prevents inflated cell counts, and improves transcript recall, ensuring that downstream analyses better reflect the true heterogeneity and organization of the tissue.

##### 1.5 Relationship between segger and existing cell segmentation methods

To offer practical advantages in accuracy, speed and usability, segger builds on a combination of existing strategies (Supp. Table B): imaging-based segmentation used by methods such as Cellpose [13] to derive an initial transcript–cell assignment based on DAPI nucleus or cell membrane staining, and modeling point-based representations of transcripts as employed by Baysor [14], Proseg[15], and ComSeg [16]. To combines these concepts, segger systematically represents transcripts, their colocalization patterns and the relationship between transcripts and cells as a heterogeneous graph that is fed to a GNN as input. While GNNs have previously been considered for cell segmentation[17], segger is the first to leverage heterogeneous graphs and to frame cell segmentation as a transcript-to-cell link prediction task. Thus, segger also builds on modern graph-based learning techniques, particularly heterogeneous multi-head graph attention networks. Beyond improving segmentation performance, the heterogeneous graph framework enables the identification of both cells and distinct cell fragments lacking nuclei, all within the same model and end-to-end training run.

segger is implemented using robust and scalable software, and is unique in offering built-in multi CPU and GPU acceleration (implemented using pytorch-geometric [18] and pytorch-lightning [19], see Appendix D), rendering it more scalable than existing methods. Speed combined with built-in heuristics to determine key model parameters are indispensable for atlas-scale cell segmentation of iST data.

#### 2 Heuristics and parameter selection guidelines

While many default parameters, such as network architecture hyperparameters, work broadly out of the box, initial graph construction and downstream prediction parameters need to be set with the properties of a given dataset in mind. We list considerations below, and provide additional instructions in the online documentation[1].

Transcript–transcript graph construction Graph parameters should be chosen to (1) maximize the correct assignment of transcripts to their cell, while (2) limiting the extent of these connections to prevent message passing between different cells, which could introduce false assignments. To help achieve this, the fixed radius used to restrict edges, `dist_tx` , should be scaled proportional to cell sizes in the sample.

When the initial segmentation is based on nuclear segmentation, `dist_tx` should be set to facilitate message passing from nuclear to cytoplasmic transcripts without extending beyond the expected cellular boundary. The number of GNN layers controls message-passing reach; we recommend 3–5 layers (corresponding to 3-5 hop neighborhoods) to avoid over-smoothing [20] (see GNN model parameters section below). Given that we typically observe cytoplasmic transcripts within 1.5–2.5 nuclear radii from the nucleus, we recommend setting `dist_tx` to half the median minimum bounding radius of nuclei in the sample (e.g., 3-hops $\times$ 0.5 radii = 1.5 radii distance).

We set `k_tx` based on transcript density within regions of size `dist_tx` . To avoid basing `k_tx` on sparsely populated slide regions, we compute densities only using transcripts located within prior segmentation boundaries (i.e., nuclei), estimating this density as $\text{density}(p_i) = \frac{A(p_i)}{n_i}$, where $A(p_i)$ is the boundary polygon's area and $n_i$ corresponds to the number of transcripts overlapping the initial polygon mask. To prevent disconnected transcripts that would hinder message passing, we err on the side of higher connectivity by selecting the 90th percentile of the density distribution:

$$\texttt{k\_tx} = \left\lceil Q^{0.9}\Big(\pi \times (\texttt{dist\_tx})^2 \times \text{density}(p_i)\Big)\right\rceil \tag{6}$$

where $Q^{0.9}$ denotes the 90th percentile across all nuclei. We note this approach also allows for adaptation of `k_tx` to varying transcript densities across different iST technologies.

##### 2.1 Tiling parameters

Tile dimensions are either set explicitly by `tile_width` and `tile_height` , or according to `tile_size` , the desired number transcripts per tile. Because entire tiles are processed during model training and inference, we recommend choosing `tile_size` according to memory constraints. For example, we chose `tile_size` =50000 to perform training on a GPU with 8GB of memory using a batch size of 3. To prevent cells from being split across tiles, adjacent regions are expanded by an overlapping margin set by a `tile_margin` parameter. We recommend setting this to the approximate maximum diameter of a cell in the units of the sample. By default, we set this to 10 (i.e., 10 µm in a Xenium sample), though this value should be adjusted based on the cell type composition of the sample.

##### 2.2 GNN model parameters

segger supports both token-based and embedding-based transcript representations. For token-based representations, `num_tx_tokens` is set to the number of genes (or slightly more) to prevent over-parameterization,

[1] https://elihei2.github.io/segger_dev

while optional scRNA-seq-derived embeddings provide additional biological insights when available. Our default model employs three mid-layers ( `num_mid_layers` = 3) to allow 3-hop information propagation without over-smoothing, with `hidden_channels` = 64 and `out_channels` = 16 to balance capacity and efficiency. Multi-head attention ( `heads` = 4) captures diverse local contexts while maintaining GPU efficiency. Training is executed with PyTorch-lightning, utilizing multi-GPU and mixed-precision support, where the number of devices is controlled by the `devices` parameter.

#### 2.3 Prediction of transcript–cell links

For the receptive field used in transcript–cell assignment, we suggest two key parameters: `k_bd` and `dist_bd` . The parameter `k_bd` , set by default to 3, limits the number of candidate cells considered per transcript. Meanwhile, `dist_bd` , with a default value of 10.0, sets the approximate cell radius to define the maximum spatial extent for candidate selection. These defaults are based on typical cell dimensions.

#### 2.4 Segger confidence score cut-off for transcript–cell link prediction

The `score_cutoff` parameter sets the minimum confidence score required to assign a transcript to a cell, based on the inner product similarity of their learned embeddings. The default threshold is 0.7, which generally favors high purity in cell assignments while maintaining sufficient transcript coverage. However, we recommend adjusting this threshold based on analytical goals or on an elbow analysis of the score distribution, which should reflect tissue heterogeneity.

#### 2.5 Transcript–transcript receptive fields for identifying fragments

To group unassigned transcripts into fragments, we build a transcript–transcript receptive-field graph using a radius-restricted $k$ NN search controlled by `k_tx` and `dist_tx` . The default `k_tx` is 4 neighbors, preserving local connectivity, and default `dist_tx` (maximum Euclidean distance for neighbor inclusion) is 5. These defaults, calibrated to typical spatial transcript densities, can be adjusted to optimize grouping accuracy, fragment size, and computational efficiency for a given dataset.

### 3 Data analysis

#### 3.1 Default 10x cell segmentation

Four our benchmarking, we consider two segmentation methods provided by 10x Genomics:

**10x Nucleus.** The Xenium Onboard Analysis software detects nuclei from DAPI images using a proprietary neural network for nucleus segmentation, which is similar to Cellpose [13]. In the default 10x Nucleus segmentation algorithm, nuclei with at least 5% of their pixel intensity above a threshold of 100 photoelectrons are retrained, and transcripts that overlap with the 2D segmentation boundaries of a given nucleus are assigned to that nucleus.

**10x Cell.** The Xenium Onboard Analysis software employs a heuristic cell boundary expansion step after nucleus segmentation to assign cytoplasmic transcripts to nuclei. Specifically, nucleus boundaries are expanded by 15 µm or until they encounter another cell boundary in X-Y space, and overlapping boundaries are resolved using an approach conceptually similar to Voronoi tessellation.

All 10x Cell and 10x Nucleus segmentations in this manuscript were generated using Xenium Onboard Analysis versions 1.0–1.9; which do not include any reported segmentation parameter changes.

#### 3.2 Breast cancer Xenium dataset

##### 3.2.1 Dataset description

We downloaded Xenium in situ spatial transcriptomics data generated from a single FFPE block of invasive breast cancer (ductal carcinoma, TNM stage T2N1M0, ER+/HER2+/PR–) from the 10x Genomics website (link). Decoded transcript data, nuclear and cellular segmentation boundaries, transcript assignments by nuclear and cellular segmentation , and DAPI images obtained from Xenium Onboard Analysis (v1.0.1) yielded ∼28,000,000 high-confidence transcripts (Xenium's quality value $\geq$ 30) and ∼168,000 segmented nuclei.

##### 3.2.2 segger

Segger default cell segmentation (i.e., with prior scRNA-seq gene embeddings) as well as tokenized (i.e., one-hot encoding) and fragment segmentation was run using default parameters and settings. Based on data-driven heuristics on the whole slide, graphs of transcript–transcript spatial neighborhoods were constructed by linking each transcript to its 20 nearest neighbors within approximately $2.5\mu$m—roughly half the median nuclear radius in the tissue. Negative edges were sampled at a 1:5 ratio to balance the graph. The slide area was partitioned into about 1,500 tiles, each containing roughly 80,000 transcripts, to enable mini-batch training on multiple 20 GB-partitioned Nvidia A100 GPUs (by default 4); these tiles were split into training, testing, and validation sets in proportions of 0.7, 0.2, and 0.1, respectively. Transcript feature vectors were computed as `gene−cell−type` abundance embeddings using a human breast cell atlas [21], with cells uniformly down-sampled to 10% and with raw counts. The default segger model was configured with an initial embedding dimension of 8, 3 intermediate hidden channel dimension of 64, and an output dimension of 16, four attention heads with sum aggregation at each layer. Training was performed for 200 epochs with a batch size of 2 for both training and validation, and predictions were generated using the same graph parameters; transcripts with assignment scores below 0.5, as determined by post-prediction score distribution analysis, were subsequently grouped into fragments. For both segger and segger-tokenized prediction of cells and fragments, transcript–cell receptive fields of $K = 4$ and $r = 5$ and transcript–transcript of $K = 5$ and $r = 3$ were considered.

##### 3.2.3 Baysor

The Xenium dataset was segmented using Baysor (v0.6.1), on julia (v1.9.2), with a regular-grid tiling strategy to handle the large computational cost of the method on a Xeon® Gold 6126 CPU. As prior segmentation, the 10x segmentation with confidence of 0.1 was applied to balance the capture of transcriptionally pure clusters with adherence to segmentation labels. Each tile was processed independently, and the resulting segmentation outputs were merged to produce a comprehensive cell segmentation map.

##### 3.2.4 BIDCell

The Xenium dataset was further processed using BidCell (bidcell==1.0.3) with the default parameters specified in the xenium.yaml configuration file from the BidCell GitHub repository (`https://github.com/SydneyBioX/BIDCell`). The analysis was executed on two Xeon® Gold 6126 CPUs (24 cores/48 threads total) and four Titan RTX GPUs (23.6 GB each). Default settings were employed for cell segmentation and transcript assignment, ensuring a standardized and reproducible workflow.

##### 3.2.5 Preprocessing of segmented cells

All datasets underwent the filtering of transcripts with Q-scores below 30, provided by the Xenium transcripts file output. For all methods other than BIDCell—which directly generates cell masks and yields a cell × gene matrix—transcripts were aggregated to their assigned cells; for 10x Nucleus, only transcripts overlapping a nucleus in the 2D projection were grouped into the corresponding cell, and an AnnData object (`anndata v0.10.9, scanpy v1.10.3`) containing a cell × gene matrix was created. Cells with fewer than 5 counts were then filtered out. For each cell, the convex hull of transcript $x, y$ coordinates was computed using `SciPy`'s ConvexHull function to estimate cell area. Data normalization was performed using `scanpy.pp.normalize_total` with `target_sum = ` $1e10^4$ followed by `scanpy.pp.log1p` for log transformation; both normalized and raw data were stored in separate layers of the AnnData objects. Cell type annotations (`"celltype_major"`) were transferred from the normalized scRNA-seq breast cancer atlas via a graph-based label transfer method (`scanpy.tl.ingest`), and UMAP embeddings on normalized counts were computed using `scanpy.tl.umap` with default parameters following neighborhood graph construction using `scanpy.tl.neighbors`.

##### 3.2.6 Cell type marker genes for benchmarking from scRNA-seq

Marker genes were identified from the breast cancer scRNA-seq atlas by performing differential expression analysis using `scanpy.tl.rank_genes_groups`, which computes, for each cell type $c$ and each gene $g$, a statistic reflecting the difference in expression between cells in $c$ and all other cells. For a given cell type $c$, the average expression of gene $g$ is computed as

$$\bar{E}_{g,c} = \frac{1}{N_c} \sum_{i \in C} E_{i,g},$$

where $C$ denotes the set of cells in $c$ and $N_c = |C|$. The distribution $\{\bar{E}_{g,c}\}_g$ is then used to define thresholds: a high-expression cutoff $T_{\text{high}}$ is determined as the $(100 - p)$th percentile (using `numpy.percentile` from `numpy v1.26.4`) of $\{\bar{E}_{g,c}\}$, and a low-expression cutoff $T_{\text{low}}$ is set as the $q$th percentile. In addition, the fraction of cells in $c$ expressing gene $g$ is calculated as

$$f_{g,c} = \frac{1}{N_c} \sum_{i \in C} \mathbb{I}\{E_{i,g} > 0\},$$

where $\mathbb{I}\{\cdot\}$ is the indicator function. Genes satisfying $\bar{E}_{g,c} \geq T_{\text{high}}$ and $f_{g,c} \geq \rho$ (with $\rho$ typically set to 0.5, meaning the gene is expressed in at least 50% of cells) are selected as positive markers, while those with $\bar{E}_{g,c} \leq T_{\text{low}}$ are designated as negative markers.

##### 3.2.7 Mutually Exclusive Co-expression Rate (MECR)

Following [22], we computed two key metrics at the transcript level to assess segmentation quality. The primary metric, MECR, quantifies the degree of mutual exclusivity between a pair of genes, defined for a given gene pair $(g_1, g_2)$ as

$$\text{MECR}(g_1, g_2) = \frac{P(g_1 \wedge g_2)}{P(g_1 \vee g_2)} = \frac{\frac{1}{N}\sum_{i=1}^{N} \mathbb{I}\{E_{i,g_1} > 0 \text{ and } E_{i,g_2} > 0\}}{\frac{1}{N}\sum_{i=1}^{N} \mathbb{I}\{E_{i,g_1} > 0 \text{ or } E_{i,g_2} > 0\}},$$

where $E_{i,g}$ denotes the expression level of gene $g$ in observation $i$, $\mathbb{I}\{\cdot\}$ is the indicator function, and $N$ is the total number of observations. A lower MECR indicates higher mutual exclusivity between the two genes.

To determine suitable gene pairs, exclusive markers were first identified from the breast cancer scRNA-seq atlas. We defined the top 30% of genes as positive markers and the bottom 5% as negative markers based on average expression, and required that a gene be expressed in at least 50% of cells within its cell type. For each cell type $c$, a gene $g$ is deemed exclusive if

$$\frac{1}{N_c} \sum_{i \in c} \mathbb{I}\{E_{i,g} > 0\} > 0.20 \quad \text{and} \quad \frac{1}{N_{\neg c}} \sum_{i \notin c} \mathbb{I}\{E_{i,g} > 0\} < 0.05,$$

where $N_c$ and $N_{\neg c}$ are the number of cells within and outside cell type $c$, respectively. The set of all mutually exclusive gene pairs are then formed by pairing exclusive markers from different cell types. This yielded 236 mutually exclusive gene pairs in the breast cancer dataset. The MECR was computed for each gene pair using the expression data from each segmentation method.

##### 3.2.8 Positive Marker Purity (PMP)

PMP quantifies the proportion of transcript counts in each cell that arise from cell-type-specific marker genes. Formally, for each cell $i$ in a given cell type $C$, let $T_i$ denote the total transcript count and $T_i^+$ denote the sum of counts for the positive markers. The PMP for cell $i$ is defined as

$$P_i = \frac{T_i^+}{T_i},$$

with $P_i$ set to zero if $T_i = 0$. This metric is computed on a per-cell basis and yields values between 0 and 1. To reduce computational overhead, when a cell type contains more than a predefined maximum number of cells (e.g., 2000), a random subset of cells is selected for analysis. Positive markers were identified by differential expression analysis of the breast cancer scRNA-seq atlas using `scanpy.tl.rank_genes_groups` and a 90% percentile threshold for high expression, and requiring that a marker is expressed in at least 50% of cells within its cell type. The resulting positive marker purity values are then aggregated by segmentation method and cell type, enabling quantitative comparisons of segmentation performance.

###### 3.2.9 Compute time and memory benchmarking

To assess the computational efficiency of segger, we measured the empirical compute time and memory requirements for segmenting the full breast cancer dataset, separately considering each of three steps of the segger workflow.

1. **Heterogenous graph construction**: Runtime and memory usage were measured for 1, 2, 4, and 8 parallel CPU cores on a system with Intel Xeon E5-2620 v4 (8-core, 2.10 GHz) and Intel Xeon E5-2660 v4 (14-core, 2.00 GHz) processors.
2. **GNN training**: Multi-GPU scalability of segger was assessed by training the model on 1, 2, and 4 NVIDIA A100 GPUs, on a system with AMD EPYC 7763 (64-core) processors. Runtime, system memory usage, and GPU memory consumption were tracked using PyTorch Lightning's `GPUStatsMonitor` callback.
3. **Downstream prediction and segmentation**: Empirical runtime and memory requirements for cell segmentation were assessed with and without fragment identification on NVIDIA A100 GPU and AMD EPYC 7763 (64-core) processors. Runtime, system memory usage, and GPU memory consumption were tracked using PyTorch Lightning's `GPUStatsMonitor` callback.

##### 3.3 NSCLC Xenium dataset

###### 3.3.1 Xenium multimodal dataset generation

FFPE blocks were generated from a resected soft tissue metastatic mass from the spine, identified as poorly differentiated carcinoma with glandular, squamous, and focal neuroendocrine features. FFPE blocks were sectioned and processed according to 10x Genomics user guidelines (CG000580, CG000582, CG000584). A customized gene panel was used for transcript detection, based on the default lung panel with 100 additional genes targeting pathways in lung cancer. Genes were selected to exclude housekeeping and highly over-expressed genes to minimize optical crowding and were finalized in collaboration with the 10x Genomics team. Decoded transcript data, nuclear and cellular segmentation boundaries, transcript assignments by nuclear and cellular segmentation, and DAPI images were obtained from the Xenium Onboard Analysis (instrument software v1.7.6.0, analysis v1.7.1.0). The dataset contains 54,641,863 transcripts and 8,603,518 segmented nuclei.

###### 3.3.2 Cell membrane staining

The slides were retrieved from the Xenium instrument and washed thrice with 500 of 0.05% PBS-T. The slides were then incubated in the blocking buffer (1X PBS pH7.4, 0.1% Tween-20, 10% heat-inactivated FBS and 10 mg/ml dextran sulfate) for 1 hr at room temperature. Slides were stained with Na-K ATPase (EP1845Y, Abcam, 1:500) conjugated with Alexa Fluor 647 overnight at 4°C. Slides were washed thrice for 10 min each and then counterstained with 500 of DAPI (5 /ml) for one min. Finally, slides were washed thrice with 500 PBS-T for one minute each and mounted using SlowFade Diamond (Invitrogen, S36963). Slides were scanned using Mirax slide scanner at 20x magnification.

###### 3.3.3 Cellpose segmentation of membrane-stained IF images

Microscopy images were converted from `MRXS` to `TIFF` using the `bioformats2raw` (v0.9.1) and `raw2ometiff` (v0.5.0) packages, generating 1024-pixel tiles with eight pyramidal levels. Images were clipped to the 1st and 99th intensity percentiles to improve contrast and reduce outlier effects, then rescaled to an 8-bit range $[0, 255]$ for compatibility with image processing tools.

Image alignment was performed in two stages to align the DAPI image from the IF dataset (moving) with the DAPI image from the Xenium dataset (fixed). First, an affine transformation was applied using the SIFT

(Scale-Invariant Feature Transform) algorithm from the `opencv-python` package (v4.6.0) to detect key points and compute a $3 \times 3$ homography matrix. This matrix was used to warp the IF DAPI and membrane images with the `warp` function from the `skimage` package (v0.21.0). For efficiency, the fourth pyramidal level (16× downsampling) was used at this stage. Nonlinear warping with `MIRAGE` (v0.0.1) was subsequently applied to refine alignment and correct local distortions. MIRAGE uses a neural field model trained on the structural similarity index (SSIM) of local patches to produce a smooth pixel-wise transformation. It was run with 196 neurons and default parameters on the first pyramidal level (2× downsampling), and the output image was used for segmentation.

Cell segmentation was performed using the `Cellpose` (v3.0.10) deep learning model with model_type = 'cyto3'. Segmentation masks were generated from channel 0 (DAPI) for nuclei detection and channel 1 (membrane) for cytoplasmic segmentation. To account for the mix of membrane-based and nuclear-based segmentations, the cell diameter was set to 0 for automatic size estimation, and the flow threshold was adjusted to 0.0 after observing a number of ill-shaped partial ROIs in the segmentation. The `cell_prob` threshold was lowered to –1 to include membrane-free nuclear masks (non-epithelial cells), which tended to have lower probabilities than membrane-stained cells (epithelial cells). Segmentation masks were mapped to transcripts by converting their $X, Y$ coordinates, scaled to image space, into pixel indices, thereby assigning each transcript to the corresponding mask label. These transcript-to-mask assignments define the Cellpose-based cell segmentation. Masks were also converted to Shapely (v2.0.1) `Polygon` objects using `skimage.measure.find_contours`, organized into a GeoSeries, and saved as GeoJSON files for visualization (**Fig. 3a,b** and **Extended Data Fig. 3.1**).

##### 3.3.4 segger

Graphs for transcript spatial neighborhoods were constructed a maximum distance cutoff dist $= 2.315\,\mu$m, corresponding to half the median nuclear radius. The value of $k$ was set to the 90th percentile of the number of neighboring transcripts within this radius rounded to the nearest integer, $k = 20$. Negative edges between boundaries and transcripts were sampled at a 1:5 ratio. Using the method described in Section 2.1.5, the slide area was divided into 1,920 tiles, each targeting approximately 100,000 transcripts, to enable mini-batch training on a 20 GB-partitioned Nvidia A100 GPU. Tiles were split into training, testing, and validation sets in proportions of 0.7, 0.2, and 0.1, respectively. Transcript feature vectors were assigned as gene–cell-type abundance embeddings. These embeddings were generated from the integrated human lung cell atlas dataset ([23]) obtained from the CellxGene Datasets portal. Cells were uniformly downsampled to 10% of the original population, and transcript counts were taken from the `SoupX-corrected` layer. The `"cell type"` column was used to assign cell type identity.

The segger model was instantiated with an initial embedding dimension of 16, a hidden channel dimension of 64 for the intermediate layer, and an output dimension of 16. One intermediate layer and four attention heads were used. Sum aggregation was applied at each layer to group transcript and boundary node embeddings. The model was trained for 100 epochs with a batch size of 2 for both training and validation. Predictions were run without connected components, using the same graph parameters as in dataset construction. An initial score cut of 0.0 was used to retain all transcripts; the score distribution was analyzed post-prediction, and transcripts with scores below the kneepoint of 0.75 were discarded.

##### 3.3.5 Baysor

Baysor was run on the same dataset using a nuclear segmentation prior from the original Xenium dataset and a prior confidence of 0.5, which balances capture of transcriptionally pure clusters with adherence to segmentation labels; in practice, lower confidence values caused over-segmentation, while higher values reduced transcript capture. Additionally, the following Baysor parameters were used: `min_molecules_per_gene = 10`, `min_molecules_per_cell = 50`, `n_clusters = 8` (based on the expected number of compartments in the dataset), and `iters = 500`. Transcript inputs were pre-filtered to exclude negative control probes and to retain those with a QV score $> 30$, consistent with the filtering criteria used for segger inputs.

Due to Baysor's extensive runtime requirements and the large dataset size, the slide area was subdivided into 12 regions with a balanced number of transcripts using segger's `get_balanced_regions` method. After processing, segmented regions were aggregated by assigning each transcript to the Baysor cell ID with the highest number of transcripts.

##### 3.3.6 Comparison to segmentation of epithelial membrane-stained images

Assignments of transcripts to segmented cells by `Cellpose` (see section 3.3.3 for more details) were transformed into an AnnData object (v0.10.9) by aggregating transcripts assigned to the same predicted cell ID and feature name, generating a cell × gene count matrix. Cells with fewer than five transcripts were removed to ensure reliable cell compartment assignment. Raw count matrices were normalized to the median library size and log-transformed (base $e$) with a pseudo-count of 0.1.

Cell compartment labels were assigned using the `CellTypist` (v1.6.3) package, transferring annotations from the integrated lung atlas scRNA-seq dataset used for segger embeddings ([23]). Fine-grained cell type annotations were grouped into five broader categories matching the compartments targeted by the Xenium gene panel: epithelial/cancer, myeloid, lymphoid, smooth muscle/pericyte, and fibroblast (**Supp. Table 1**). To emulate transcript capture rates observed in Xenium datasets, genes not in the panel were removed from the scRNA-seq dataset and cell counts were downsampled to 100 transcripts per cell. The resulting count matrix was then log-transformed for compatibility with `CellTypist`.

To address class imbalance and preserve rare cell types, `CellTypist`'s guidelines for large atlas label transfer were followed: each cell type was downsampled to 2,000 cells with replacement, duplicates were removed, and a total of $X$ cells was retained. The model was trained with default parameters to predict cell compartment labels and applied to the `Cellpose` count matrix, which was downsampled to 100 transcripts per cell to match the training dataset and log-transformed. Cells annotated as epithelial/cancer were retained for benchmarking (**Fig. 3** and **Extended Data Fig. 3.1**).

For benchmarking comparisons, a different approach to cell type labeling was applied to avoid biases introduced by poor segmentation quality. Specifically, for segger, Baysor, and 10x segmentations, all segmented cells that shared more than one-third of their total transcripts with epithelial cells identified by `Cellpose` were compared. This approach ensured that cells mislabeled due to contamination or filtered for low UMI counts due to over-segmentation were not excluded from the comparison.

We define the following metrics for comparing each segmentation method to `Cellpose` segmentation:

- **Cell Area:** Calculated as the convex hull of transcripts assigned to each cell, using the `area` property of the `ConvexHull` class in `scipy` (v1.10.1).

- **Cell Overlap:** The number of cell IDs from alternative segmentation methods (e.g., segger, Baysor, or 10x) found within transcripts assigned to the same Cellpose-segmented cell. Only overlaps of at least five transcripts that represent at least 25% of the total transcripts in the overlapping cell are included to reduce noise.

- **Cell Contamination:** The fraction of transcripts assigned to epithelial cells by an alternative segmentation method that were not assigned to epithelial cells by Cellpose. Mathematically, for a given cell,

$$\text{Contamination} = \frac{|T_{\text{alt}} \setminus T_{\text{CP}}|}{|T_{\text{alt}}|},$$

  where $T_{\text{alt}}$ is the set of transcripts assigned by the alternative method (e.g., segger, Baysor, or 10x) and $T_{\text{CP}}$ is the set assigned by Cellpose. Cells with fewer than five transcripts are excluded.

- **Cell Recall:** The fraction of transcripts assigned to epithelial cells by `Cellpose` that are also assigned to the corresponding cell by an alternative method. This is expressed as

$$\text{Recall} = \frac{|T_{\text{alt}} \cap T_{\text{CP}}|}{|T_{\text{CP}}|}.$$

  As with contamination, cells with fewer than five transcripts are excluded from the calculation.

###### 3.3.7 Comparison of over-segmentation in segger vs. Baysor

For both segmentation methods, transcript assignments were transformed into AnnData objects (v0.10.9) and normalized as described above. The same `CellTypist` model used to annotate the `Cellpose` dataset was applied to assign cell compartment labels. Additionally, entropy across label probabilities was computed for each cell to quantify the uncertainty of cell type assignments.

For visualization of cell compartments in segger and Baysor datasets (**Extended Data Fig. 3.2c,d**), principal component analysis (PCA) was performed on log-normalized count matrices, retaining the principal components that explained 75% of the variance. Projections were generated using the UMAP implementation in `CUML` (v24.08.00) with parameters min_dist = 0.25, *n_neighbors* = 20, *n_epochs* = 1000, and default settings otherwise. Cells were pseudo-colored on a continuous gradient from gray to their assigned cell type color according to entropy values: low-entropy (pure) cells appear in the color of their assigned type, while high-entropy (mixed) cells appear gray.

###### 3.3.8 Differential expression analysis in segger vs. 10x

Differential expression analysis was performed on Epithelial cells labeled as Epithelial/Cancer (using transferred labels from `Cellpose` as described above) from segger and 10x Cell segmentations using the `pyDESeq2` (v0.4.12) package. The analysis was run on raw counts with a design factor corresponding to the segmentation method (segger or 10x Cell). To account for spatial heterogeneity in segmentation (e.g., differences in tumor histology or cell type composition affecting the success of 10x's nuclear expansion), the slide was subdivided into 12 equally sized regions, with each region treated as a replicate for $p$-value calculation. This approach avoided pseudo-bulking the entire sample, which could suppress spatial differences. In the resulting volcano plot, individual genes were colored according to the cell compartment label with the highest coefficient in the `CellTypist` model used for cell typing.

#### 3.4 Colon Xenium dataset

##### 3.4.1 Data acquisition and segmentation

Publicly available Xenium outputs for the human non-diseased colon preview dataset with pre-designed and add-on panel, as well as a post-Xenium H&E image, were obtained from the 10x Genomics website (link). The 325-gene Xenium Human Colon Panel (10x Genomics)used to profile the sample was designed to identify absorptive enterocytes, enteroendocrine cells, goblet cells, Paneth cells, fibroblasts, and vascular endothelial cells. It was supplemented with 100 additional genes, including those involved in signaling, chemokines, and stromal cell markers. The dataset was processed with Xenium Onboard Analysis (analysis v1.6.0.7) and contains 32,106,157 transcripts and 275,822 segmented cells.

Cells were segmented using segger following the approach described in Section 3.3.4, with adjustments to account for the smaller dataset size and different transcript densities. Specifically, updated parameters included `tile_size = 50,000`, `dist_tx = X`, `k_tx = 20`, `batch_size = 3`, `init_emb = 8`, `hidden_channels = 32`, `out_channels = 8`, and `epochs = 80`. Feature vectors were generated as gene–cell-type abundance embeddings from the single-cell atlas of the human intestinal tract [24], subset to samples from the adult colon, obtained from the Cell × Gene Datasets portal. Cell type annotations were taken from the column *cell_type* and counts from *.X*. Otherwise, dataset construction, training, and prediction steps remained unchanged.

##### 3.4.2 Dataset preprocessing and cell typing

All segmentations (segger, 10x Cell, and 10x Nucleus) were transformed into `AnnData` objects (v0.10.9) as described previously, with X, Y coordinates assigned to each cell based on the mean transcript coordinates. For downstream analysis, we aimed to compare segmentations in a realistic analysis scenario. Given the inherent sparsity of Xenium data, determining a suitable transcript threshold is critical: too few transcripts introduce sparsity, while filtering out cells with many transcripts risks removing meaningful populations. We filtered the segger dataset to retain cells with at least 25 transcripts, based on the knee-point in the transcript distribution, and used the same threshold for the other datasets. This threshold is consistent with similar published Xenium analyses ([25]).

For all datasets (segger, 10x Cell, and 10x Nucleus), PCA was performed on log-normalized count matrices, retaining principal components that explained 95% of the variance to enable finer cell type resolution, despite admitting more noise from lowly expressed genes. Projections were generated using UMAP (CUML v24.08.00) with the following parameters: `min_dist = 0.2`, `n_neighbors = 20`, `n_epochs = 1000`, and `local_connectivity = 2`. For clustering, we applied a custom GPU implementation of the PhenoGraph algorithm (see 'Code Availability') with `resolution = 4` during Louvain clustering.

We found that transferring cell type labels from scRNA-seq data to Xenium using CellTypist worked well for broad compartment-based annotations in the NSCLC example but struggled with fine-grained cell type assignments in our Xenium colon dataset. This limitation likely reflects differences in gene capture between the two technologies and the absence of key marker genes in the standard Xenium human colon panel compared to the much larger set of genes detectable by scRNA-seq in the intestinal atlas[24]. To address this, we compiled three curated cell type gene sets from different sources. First, we included the differentially expressed genes (DEGs) reported for each annotated cell type in the colon single-cell atlas. Second, we incorporated DEGs from the single-cell atlas of immune dysregulation in Crohn's Disease in the human colon to help resolve immune subsets and disease-related states ([26]). Third, we used the panel annotations provided by 10x Genomics. These cell type gene sets were consolidated to resolve differences in cell type naming across sources (original and re-labeled cell types are listed in **Supplementary Table 3**), subset to genes available in the Xenium panel, and marker sets without any genes following subsetting were excluded (**Supplementary Table 3**).

We began by annotating the 10x Nucleus dataset, presumed to be the least contaminated. Annotation was performed per cluster by z-scoring the log-normalized expression data and running the `score_genes` function in scanpy (v1.10.2), assigning each cluster the cell type with the highest average marker set score. Using these preliminary annotations, we trained a `CellTypist` model on the labeled 10x Nucleus dataset and annotated the segger and 10x Cell segmentation datasets using the trained model. During this process, we identified two segger clusters originally labeled as "monocytes" that were highly underrepresented in the 10x Nucleus dataset used to train the `CellTypist` model (10% overlap), encouraging us to reevaluate their cell type annotation. Differential expression analysis revealed high expression of *IL1B*, *CXCR1*, and *CXCR2*, suggesting neutrophil identity (**Extended Data Fig. 4.1a**). This was confirmed by a colon pathologist through H&E examination, and we re-annotated these clusters as "neutrophil" (**Extended Data Fig. 4.1b**).

##### 3.4.3 Hotspot analysis of neutrophils and epithelial neighbors

We used Hotspot (v1.1.2, [27]) to identify context-specific gene modules in single-cell and spatial transcriptomics datasets. Briefly, Hotspot calculates pairwise local correlations between genes in local cell neighborhoods on a user-provided cell–cell similarity graph (e.g., spatial kNN), identifying genes with high local autocorrelation. The resulting gene–gene correlation matrix is clustered to output a set of gene modules. We ran Hotspot on our segger dataset using raw counts, the *danb* model, and 20 nearest neighbors, calculated based on the spatial positions of cells. Genes with an FDR $< 0.01$ were retained for downstream analysis, resulting in 421 genes used for module discovery. Modules were identified using the `create_modules` function with default parameters (`minimum_gene_threshold = 3` and `core_only = False`), yielding 39 co-varying gene modules (**Supplementary Table 4**). We note that the lower minimum gene threshold compared to scRNA-seq is consistent with Xenium data, which has far fewer genes overall. Module scores were calculated using the `calculate_module_scores` function.

Among the 39 modules, we identified two—Module 16 and Module 26—that are significantly correlated with the distance to the nearest neutrophil cell ($R = -0.400$ and $-0.243$; $p = 0$, respectively, by Pearson correlation). Distances were computed for up to 500 nearest neighbors (cells with no neutrophil found within that range were excluded). Within epithelial subsets, upper crypt epithelial cells were defined as those with Module 3 scores above the 75th percentile. Among Module 3+ cells, we further defined a Module 16+/26+ subset as cells scoring for both modules above the 90th percentile; cells positive for Module 3 not meeting these criteria were labeled Module 16−/26−. In our analysis of neighborhood compositions for these subsets, for each cell we calculated the fraction of cells labeled as neutrophil among its 200 nearest spatial neighbors. Nearest neighbors were calculated using the `NearestNeighbors` class in `CUML` (v24.08.00) and the X, Y coordinates assigned to each cell.

To estimate the number of cell type markers in the Xenium gene panel, we re-annotated cell types in the intestinal single-cell dataset to align with our cell type definitions (**Supplementary Table 3**). After downsampling to 1,000 cells per cell type, we performed differential expression analysis using `scanpy.tl.rank_genes_groups` (v1.10.2) and retained genes with $p < 1 \times 10^{-3}$ and $\log_2 \mathrm{FC} > 3$. The number of markers for each cell type was defined as the intersection of these differentially expressed genes and those present in the Xenium panel.

##### 3.4.4 Differential expression in tuft cells

We performed differential expression analysis on tuft cells from segger and 10x Cell segmentations using the `pyDESeq2` package (v0.4.12). The analysis was conducted on raw counts with the design factor set to the segmentation method (segger or 10x Cell).

### Appendix

### A Graph Neural Networks

#### A.1 Overview

Graph Neural Networks (GNNs) are designed to operate on graph-structured data, where data is represented as nodes (entities) and edges (relationships) connecting those nodes. GNNs iteratively aggregate and transform node features by leveraging information from their neighboring nodes. This makes them effective for node-level, link-level, and graph-level prediction tasks, including both classification and regression [5, 28, 18].

##### A.1.1 Message Passing Mechanism

The core idea of GNNs is to iteratively update node representations by aggregating information from their neighbors. This process is called the message passing paradigm, where each node receives messages from its neighbors, aggregates them, and updates its own state. The general form of a GNN layer is expressed as:

$$h_i^{(l+1)} = \text{UPDATE}\left(h_i^{(l)}, \text{AGGREGATE}\left(\{h_j^{(l)} : j \in \mathcal{N}(i)\}\right)\right) \tag{7}$$

Here, $h_i^{(l)}$ represents the feature vector of node $i$ at layer $l$, and $\mathcal{N}(i)$ denotes the set of neighbors of node $i$. The **AGGREGATE** function collects information from the neighbors, while the **UPDATE** function combines this aggregated information with the current state of the node. For instance, the AGGREGATE function could be a simple sum of the neighboring node features, while the UPDATE function might apply a learnable linear transformation followed by a non-linear activation function such as ReLU. These functions are customizable and can vary based on the specific GNN architecture.

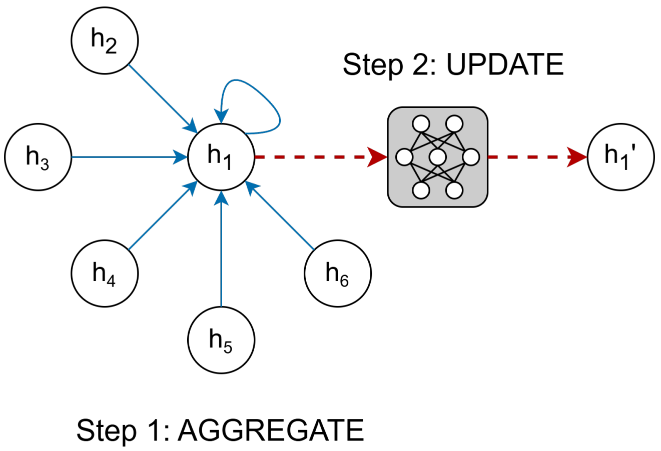

Figure 1: Message passing mechanism in Graph Neural Networks. Step 1 involves aggregating feature vectors from the neighborhood of node 1 (including its own feature vector) using a permutation-invariant function, such as summation. In Step 2, the aggregated feature vector is transformed using a learnable function, such as a Multi-Layer Perceptron (MLP), to produce the updated feature vector for node 1.

##### A.1.2 GNN Example: Graph Isomorphism Network

Graph Isomorphism Networks (GINs) utilize a simple yet expressive GNN layer : it performs a sum over the features of neighboring nodes [29]. This aggregation treats all neighbors equally—it is not a weighted sum, meaning that each neighbor's contribution is given the same importance during aggregation. The aggregated features are then combined with the node's own features through an arbitrary Multi-Layer Perceptron (MLP) as the update function. This operation can be expressed as:

$$h_i^{(l+1)} = \text{MLP}\left(\sum_{j \in \mathcal{N}(i) \cup \{i\}} h_j^{(l)}\right) \tag{8}$$

#### A.2 Graph Attention

Graph Attention Networks (GATs) extend the standard message-passing paradigm in GNNs by incorporating an attention mechanism that enables nodes to weigh the importance of their neighbors differently [4]. Unlike simpler aggregation schemes where all neighbors contribute equally to a node's representation, GATs assign attention scores to each neighboring node, allowing the model to focus more on the most relevant neighbors during the aggregation process. This makes GATs particularly powerful for tasks where the relative importance of neighbors varies significantly across different nodes. Specifically, we focus on GATv2 [6], a newer version of GAT that improves the expressiveness of the attention mechanism.

##### A.2.1 Attention Mechanism in GATs

The core innovation in GATs lies in their learnable attention coefficients, computed for each edge connecting a node $i$ to its neighbor $j$. The attention coefficient $\alpha_{ij}$ is calculated as follows:

$$e_{ij} = a^T \text{LeakyReLU}([Wh_i || Wh_j]) \tag{9}$$

where $h_i$ and $h_j$ are the feature vectors of nodes $i$ and $j$, $W$ is a learnable weight matrix that transforms the node features, and $a$ is a learnable weight vector applied to the concatenated features. Next, the attention coefficient $\alpha_{ij}$ is obtained by normalizing $e_{ij}$ across all neighbors of node $i$ using the softmax function:

$$\alpha_{ij} = \text{softmax}(e_{ij}) = \frac{\exp(e_{ij})}{\sum_{k \in \mathcal{N}_i} \exp(e_{ij})} \tag{10}$$

The resulting attention coefficient $\alpha_{ij}$ quantifies the importance of node $j$'s features when updating node $i$. Higher values indicate greater importance, allowing the model to prioritize influential neighbors while diminishing the impact of less relevant ones.

##### A.2.2 Aggregation and Update Functions

In Graph Attention Networks (GATs), the aggregation and update process for node $i$ is described by the following formula:

$$h_i^{(l+1)} = \sigma\left(\sum_{j \in \mathcal{N}_i \cup \{i\}} \alpha_{ij}^{(l)} W h_j^{(l)}\right) \tag{11}$$

- **Aggregation**: The aggregation function is the weighted sum of the features of neighboring nodes (including the node itself), where the weights are the attention coefficients $\alpha_{ij}$.
- **Update**: The result of the aggregation is passed through the update function, typically a non-linear activation such as ReLU, though it can also be more complex, like an MLP.

To increase the expressive power of the model, GATs typically use multi-head attention. In this setting, multiple independent attention mechanisms (i.e. attention heads) operate in parallel, producing multiple sets of attention-weighted representations. The outputs from the heads are concatenated:

$$h_i^{(l+1)} = \Big\Vert_{k=1}^{K} \sigma\left(\sum_{j \in \mathcal{N}_i \cup \{i\}} \alpha_{ij}^{(l,k)} W^{(k)} h_j^{(l)}\right) \tag{12}$$

where $K$ is the number of attention heads. This approach allows GATs to capture different views of neighbor importance, resulting in richer and more robust node embeddings.

#### A.3 Heterogeneous Graph Neural Networks

Heterogeneous Graph Neural Networks (HGNNs) are designed to manage graphs with diverse node and edge types, capturing complex interactions and relationships. Unlike homogeneous graphs, where all nodes and edges share the same type, heterogeneous graphs require differentiated treatment of node and edge types during message passing [8, 30]. This differentiation allows HGNNs to capture the rich interactions present in complex systems such as knowledge graphs [31, 32], recommender systems [33, 34, 35], and biomedical data [36, 37].

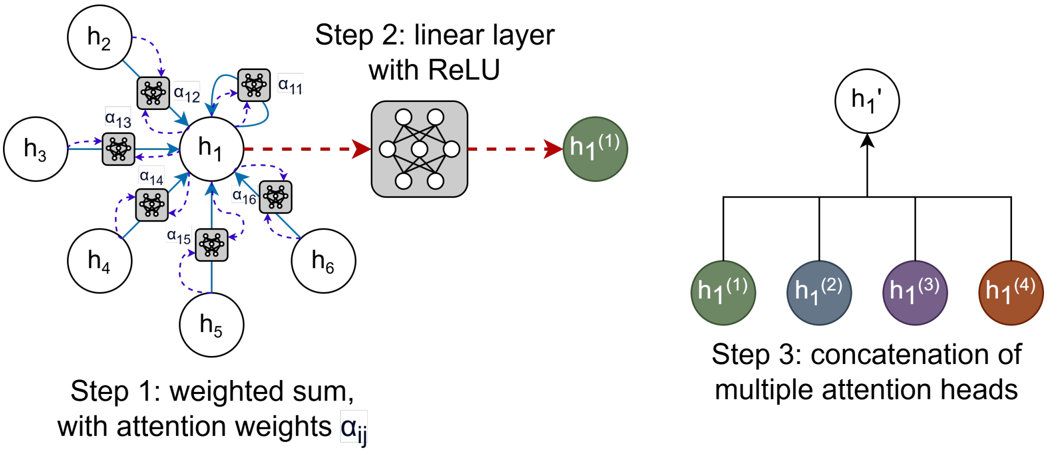

Figure 2: Graph Attention Networks (GATs) with multi-head attention. In Step 1, feature vectors from the neighborhood of node 1 $h_1, h_2, ...h_6$ are aggregated using attention weights $\alpha_{11}, \alpha_{12}, ..., \alpha_{16}$, which quantify the importance of each neighbor's feature vector. In Step 2, the aggregated features are updated through a learnable linear transformation with ReLU activation, resulting in the new feature vector $h'_1$. Optionally, in Step 3, multiple attention heads may operate in parallel, producing independent outputs that are concatenated to create a more expressive representation.

##### A.3.1 Characteristics and Challenges of Heterogeneous Graphs

Heterogeneous graphs introduce several complexities, including:

- **Multiple Node and Edge Types**: Nodes and edges represent different entities and relationships, requiring models to distinguish between them during message passing.
- **Complex Interaction Patterns**: The relationships between nodes vary in meaning and structure, requiring specialized aggregation and update functions to model these relationships.
- **Data Imbalance**: Certain node or edge types may dominate the graph, potentially leading to biased representations.

##### A.3.2 Heterogeneous GNN Layers

In HGNNs, heterogeneity is handled by defining distinct edge types that control how information flows between different node types. Each node type aggregates messages from its neighbors based on the types of nodes and edges it connects with. This approach ensures that the characteristics of each relationship are captured, allowing the model to learn their collective influence on node features.

Formally, let $R$ denote the set of edge types, and let $\mathcal{N}^{(r)}(i)$ represent the neighbors of node $i$ under relation $r$. The feature update for node $i$ at layer $l$ in an HGNN is expressed as:

$$h_i^{(l+1)} = \text{UPDATE}\left(h_i^{(l)}, \bigoplus_{r \in \mathcal{R}} \text{AGGREGATE}_r\left(\{h_j^{(l)} : j \in \mathcal{N}^{(r)}(i)\}\right)\right) \tag{13}$$

Here, $\text{AGGREGATE}_r$ denotes the relation-specific aggregation function that combines features from neighbors of node $i$ connected via relation $r$, and $\bigoplus$ denotes an outer aggregation operation that combines results across different relations.

##### A.3.3 Heterogeneous GAT with Sum Aggregation

In heterogeneous Graph Attention Networks (GATs), the standard message-passing mechanism adapts to include attention weights, which determine the importance of each neighbor's contribution based on the specific relation type. This approach allows nodes to selectively focus on the most relevant neighbors while capturing complex interactions across multiple relations.

The feature update for node $i$ at layer $l$ in a heterogeneous GAT with sum aggregation is defined as:

$$h_i^{(l+1)} = \sigma\left(\sum_{r \in \mathcal{R}} \sum_{j \in \mathcal{N}_i^{(r)} \cup \{i\}} \alpha_{ij}^{(r,l)} W^{(r)} h_j^{(l)}\right) \tag{14}$$

This formulation combines attention-based weighting with relation-specific transformations, allowing for expressive modeling of heterogeneous graphs by aggregating contributions across all types of relations.

segger also leverages a heterogeneous GAT with sum aggregation to capture complex interactions between different node types. In segger, there are two types of nodes: transcripts and boundaries, linked by two types of edges: transcript-neighbors-transcript (connecting transcripts in close spatial proximity) and transcript-belongs-boundary (indicating that the spatial location of a transcript lies within the boundary).

In this setup, the features of boundary-type nodes are updated by aggregating information from themselves and from transcript-type nodes that belong to them (via the transcript-belongs-boundary relation). Conversely, transcript-type nodes update their features by aggregating information from themselves, neighboring transcripts (via the transcript-neighbors-transcript relation), and the boundary-type node they belong to (via the transcript-belongs-boundary relation). This aggregation scheme allows segger to effectively capture the interactions between different biological entities.

#### A.4 Link Prediction with Graph Neural Networks

##### A.4.1 Problem Definition

Link prediction is a core task in graph learning, focused on determining the existence or probability of an edge (link) between two nodes in a graph. Given a graph with observed nodes and edges, the goal is to predict potential or missing links, offering insights into the underlying structure of the data and uncovering hidden relationships [5, 38].

Many real-world problems can be framed as link prediction tasks on heterogeneous graphs:

- **Knowledge Graph Completion**: Predicting missing relationships between entities (nodes) of different types, thereby enhancing the completeness and utility of knowledge bases [31, 32].
- **Recommender Systems**: Modeling interactions between users, items, and contextual factors, capturing complex relationships to deliver personalized recommendations [33, 34, 35].
- **Biomedical Networks**: Identifying interactions between diverse biological entities, such as genes, proteins, and diseases, to facilitate discoveries in health and medicine [36, 37].

Despite its broad utility, link prediction presents several challenges:

- **Scalability**: Many real-world graphs contain millions or billions of nodes and edges, requiring efficient algorithms that can scale to such massive datasets [39].
- **Sparsity**: In many graphs, only a small fraction of all possible links exist, making it challenging to identify meaningful connections among a large number of potential links [40].
- **Heterogeneity**: Real-world graphs often contain diverse node and edge types, requiring models that can capture complex interactions and relationships across different entities [41].

  Graph Neural Networks (GNNs) offer a promising solution to link prediction challenges by learning expressive node and edge representations. By capturing both graph structure and heterogeneous node/edge attributes, GNNs provide a data-driven approach for uncovering hidden relationships in complex graphs.

##### A.4.2 Using GNNs for Link Prediction

Graph Neural Networks (GNNs) are powerful tools for link prediction due to their ability to work with both node features and the underlying graph structure [5, 38]. By capturing local neighborhood connectivity and the global topology, GNNs learn expressive node representations that enable the identification of potential or missing links [9, 42]. Unlike traditional methods that rely on hand-crafted features or shallow embeddings, GNNs offer a flexible, end-to-end learning framework that adapts to various graph structures and link prediction tasks.

The typical process for link prediction with GNNs involves several steps [43, 44, 32]:

1. **Node Embedding Generation**: The GNN processes the graph to generate embeddings for each node by aggregating information from its neighbors. These embeddings capture both the node's features and its structural context.
2. **Combining Node Embeddings for Link Prediction**: To predict a potential link between two nodes, their embeddings are combined using techniques such as concatenation, element-wise multiplication, dot products, or more complex functions like Multi-Layer Perceptrons (MLPs). This combined representation is then used to score the probability of a link between the nodes.

3. **Framing Link Prediction as Binary Classification**: Link prediction is commonly framed as a binary classification task, where the goal is to predict whether an edge exists between a given pair of nodes. Positive examples correspond to observed edges in the graph, while negative examples represent node pairs without an observed edge. A common practice is to sample negative edges by selecting random node pairs that are not connected in the observed graph. During training, binary cross-entropy loss is used to minimize the difference between the predicted link probabilities and the ground-truth labels (1 for positive links, 0 for negative links).

4. **Train-Validation-Test Splits**: Like other machine learning tasks, link prediction requires careful splitting of the data into training, validation, and test sets. Typically, edges are split into these sets, while ensuring that no information from the test set leaks into training or validation. This split ensures that models are evaluated on their ability to generalize to unseen data. For negative sampling, an equivalent number of non-existent edges are often included in each split to balance the data.

This approach offers several advantages:

1. **Scalability**: By focusing on generating node embeddings, GNN-based link prediction methods can efficiently handle large graphs without explicitly modeling every possible node pair [43].

2. **Efficiency**: Once learned, node embeddings can be reused for different tasks, enabling fast and flexible link predictions [45].

Predicting links in heterogeneous graphs is more complex due to the presence of different node and edge types. Unlike homogeneous graphs, where all nodes and edges are treated the same, heterogeneous GNNs use relation-specific transformations and aggregation strategies [31]. This means that interactions between different types of nodes and edges are processed uniquely, allowing the model to capture specific patterns for each type.

##### A.4.3 Evaluation Metrics for Link Prediction

Evaluating link prediction performance is crucial to assess how well a model predicts the existence of links between nodes. Commonly used metrics include:

- *Accuracy*: Measures the proportion of correctly predicted links (both positive and negative) out of the total predictions made. While accuracy is a straightforward measure, it may not always be the best metric for imbalanced datasets, where negative examples (non-existent links) can far outnumber positive examples (existing links).

- *AUROC (Area Under the Receiver Operating Characteristic Curve)*: This metric evaluates the model's ability to distinguish between positive and negative links across different decision thresholds. A higher AUROC value indicates better discrimination, with a value of 1 indicating perfect distinction and 0.5 indicating random guessing.

- *Precision, Recall, and F1-Score*: These metrics provide a more nuanced view of link prediction performance:
  - *Precision*: The proportion of correctly predicted positive links out of all predicted positive links. High precision indicates a low rate of false positives.
  - *Recall*: The proportion of correctly predicted positive links out of all actual positive links. High recall indicates a low rate of false negatives.
  - *F1-Score*: The harmonic mean of precision and recall, providing a balanced measure that accounts for both false positives and false negatives. It is useful when there is an uneven class distribution.

When evaluating link prediction models, it is also important to compare their performance against appropriate baselines. Baselines may include:

- *Random Link Prediction*: Predicting links randomly, serving as a naive benchmark.

- *Traditional Heuristics*: Metrics such as common neighbors, preferential attachment, and Jaccard coefficient, which are simple but effective graph-based methods used historically for link prediction.

### B Comparison of segger and existing segmentation approaches

Table 1: Comprehensive comparison of cell segmentation methods for spatial transcriptomics. The table summarizes key attributes including overall model architecture (categorized as Image based (CNN), Probabilistic, Graph-Based, Staining-Free, or Point-density based), utilization of transcript co-localization information, requirements for nucleus (DAPI) and membrane staining (Required, Optional, or Not used), incorporation of morphological features, and the role of scRNAseq data (Required, Optional, or Not used). It also details computational speed (in hours) and software availability.

| **Method** | **Overall Model** | **Tx-co-localization Info** | **Nucleus (DAPI) Staining** | **Membrane Staining** | **Morphological Features** | **scRNAseq Info** | **Speed (hours)** | **Software** |
|---|---|---|---|---|---|---|---|---|
| Nuclei Expansion (Voronoi diagram) [46] | Point-density based (Voronoi) | No | Required | Not used | No | No | Fast (<5 h) | Widely available, simple to implement |
| **segger** | Graph-Based (GNN: link prediction) | Yes | Optional | Optional | Yes | Optional | Fast (<5 h) | GitHub; CLI/API docs; tutorials; built-in multi CPU/GPU acceleration |
| Baysor[14] | Probabilistic (Bayesian + MRF) | Yes | Optional | Optional | No | No | Slow (>24 h) | GitHub (Julia/Docker); well-documented |
| BIDCell[47] | Image based (CNN: U-Net3+) | Yes | Required | Not used | Yes | Required | Fast (<5 h) | GitHub; well-documented; tutorials |
| Cellpose[13] | Image based (CNN) | No | Optional | Optional | Yes | No | Fast (<5 h) | Python/GUI; well-documented |
| ProSeg[15] | Probabilistic (simulated annealing) | Yes | Optional | Optional | Yes (if using image data) | No | Moderate (5–24 h) | GitHub (C++/Rust); limited docs |
| PCiSeq[48] | Probabilistic | Yes | Required | Not used | Yes | Optional | Fast (<5 h) | GitHub; well-documented; tutorials |
| Mesmer[49] | Image based (CNN) | No | Required | Optional | Yes | No | Fast (<5 h) | GitHub; well-documented; tutorials |
| Omnipose[50] | Image based (CNN: gradient segmentation) | No | Required | Optional | Yes | No | Fast (<5 h) | GitHub; well-documented; tutorials |
| SAM[51] | Image based (Transformer) | No | Optional | Optional | Yes | No | Fast (<5 h) | Pre-trained model + docs; custom prompts |
| ComSeg[16] | Graph-Based (graph clustering) | Yes | Optional | Optional | No | No | Moderate (5–24 h) | GitHub; well-documented; tutorials |
| GeneSegNet[52] | Image based (CNN: gene-integrated) | Yes | Required | Not used | Yes | No | Slow (>24 h) | GitHub; limited documentation |
| JSTA[53] | Point-density based (Watershed) | Yes | Required | Not used | No | Required | Slow (>24 h) | GitHub (R/Python); lab-provided |
| Bering[17] | Graph-Based (GNN: adjacency) | Yes | Required | Not used | No | No | Slow (>24 h) | GitHub; limited docs |
| UCS[54] | Image based (CNN) | Yes | Required | Not used | Yes | No | Slow (>24 h) | GitHub; limited docs |
| Ilastik[55] | Image based (interactive ML) | No | Optional | Optional | Defined by user | No | Fast (<5 h) | Widely available; well-documented; GUI-based |
| SCS[56] | Image based (Transformer) | Yes | Required | Not used | Yes | No | Slow (>24 h) | Authors' code; Requires custom implementation |
| SSAM[57] | Staining-Free | Yes | Not used | Not used | No | Optional | Fast (<5 h) | GitHub; well-documented |
| Watershed (Nuclei)[58] | Point-density based (Watershed) | No | Required | Optional | No | No | Fast (<5 h) | Widely available (ImageJ, skimage) |

### C *segger's* implementation and workflow

#### C.1 Data preprocessing

The segger software comes with a preprocessing pipeline to transform spatial transcriptomics data into a structured format suitable for cell segmentation, formulated as a link prediction task. The input data consists of transcripts with spatial coordinates and nucleus or cell type boundaries. The spatial data is divided into smaller regions called tiles, with each tile corresponding to a graph representation. This tile-based approach preserves local spatial relationships and enables detailed modeling of interactions within each region. These graphs are then saved as *PyTorch Geometric* (*PyG*) data objects and are randomly allocated into training, validation, and test sets for model training and evaluation.

##### C.1.1 Input data

The data preprocessing for segger begins with a set of required input files.

- `transcripts.parquet` : Contains spatial and metadata information for individual transcripts.
  - **Spatial coordinates**: Indicate the positions of transcripts within the spatial sample.
  - **Transcript ID**: A unique identifier for each transcript.
  - **Feature label**: Represents the gene associated with each transcript.
  - **Overlap status**: Specifies whether a transcript spatially overlaps with a nucleus/cell boundary.
  - **Cell ID**: Links the transcript to a specific nucleus/cell, if applicable.
  - **Quality value**: Reflects the measurement confidence of the transcript.
- `nucleus_boundaries.parquet` or `boundaries.parquet` : Contains boundary and metadata information for either for nuclei or cells.
  - ***Node coordinates***: Define the spatial boundary of the nucleus/cell.
  - ***Cell ID***: A unique identifier for each nucleus/cell.

***Optional enrichment***: If single-cell RNA sequencing (scRNA-seq) data is available, it injects biological prior knowledge through gene-cell-type abundance embeddings, typically enhancing model performance. If not specified, transcripts are represented by one-hot encodings based on their feature labels.

##### C.1.2 Configuration settings

Preprocessing is guided by a sample-specific YAML configuration file. For Xenium datasets, two example configurations are provided: `xenium.yaml` for nucleus boundaries ( `nucleus_boundaries.parquet` ) and `xenium_v2.yaml` for cell boundaries ( `boundaries.parquet` ).

##### C.1.3 Required parameters

Several parameters guide the data preprocessing workflow, either specified through the CLI or a YAML configuration file:

- `base_dir` : Base directory containing the raw input files.
- `data_dir` : Directory for saving the processed segger dataset.
- `sample_type` : Specifies the sample type (e.g., xenium, merscope), determining the YAML configuration to use.
- `scrnaseq_file (optional)` : Path to an scRNA-seq file (anndata object saved in `.h5ad` format) for computing gene-cell-type abundance embeddings.
- `celltype_column (optional)` : Column in the scRNA-seq file (in `.obs` of the `anndata` object) containing the cell-type annotations.
- `k_tx` : Number of nearest neighbors to consider for transcript nodes.
- `dist_tx` : Maximum distance for connecting transcript nodes.
- `tile_width/tile_height` : Dimensions of spatial tiles.
- `neg_sampling_ratio` : Ratio of negative edges to positive edges, sampled for training.
- `val_prob/test_prob` : Proportion of data allocated for validation/testing.
- `n_workers` : Number of parallel workers for data processing.

##### C.1.4 Initialization

Initialization involves the following key steps:

1. **Loading YAML configuration and metadata extraction**
   - The YAML file corresponding to the specified sample type is loaded to guide data processing. Metadata is then extracted from the input files.
   - **Transcripts metadata**: Extracted from `transcripts.parquet` and includes spatial coordinates, transcript IDs, feature labels, overlap status, associated cell IDs, and quality values. A list of substrings used for filtering unwanted features (e.g., control probes) is gathered. If a single-cell RNA sequencing (scRNA-seq) file is provided, any genes present in the transcripts but missing from the gene-cell-type embeddings are added to this list of filter substrings.
   - **Boundaries metadata**: Extracted from `nucleus_boundaries.parquet` or `boundaries.parquet` and includes node coordinates and associated cell IDs.
2. **Embedding initialization**
   - If a scRNA-seq file is specified, gene-cell-type abundance embeddings are computed to enrich transcript features. This calculates the percentage of cells within each cell type that express each gene, creating a matrix where rows represent genes and columns represent cell types.
   - Transcripts are then enriched with these gene-cell-type embeddings if available. If no scRNA-seq data is provided, transcripts are instead represented using one-hot encodings of their feature labels.
3. **Setting up data storage structure**: The data directory is prepared to store processed tiles. This includes creating separate subdirectories for training, validation, and test tiles. Each generated tile is randomly assigned to one of these sets.

##### C.1.5 Balanced region generation

To enable efficient parallel processing, following initialization, the spatial transcriptomics sample is divided into smaller, balanced regions, preserving local spatial relationships within each region. The main points are the following:

1. **Generating balanced regions using an ND-Tree structure**: To achieve balanced partitioning and efficient parallel processing, the spatial extent of the whole sample is divided into smaller regions using an ND-Tree data structure. This approach relies on the spatial distribution of boundaries, which typically forms a smaller, well-defined subset compared to transcripts, while still representing the overall spatial layout.
   - **Initialization and recursive splitting**: The entire area covered by the sample is initially represented as a bounding rectangle, defined by the minimum and maximum spatial coordinates of the boundaries. The ND-Tree recursively splits the sample along the dimension with the largest spatial spread to maintain balanced partitioning. Each split creates two child nodes, distributing data points based on their position relative to the split point. This process continues until each region contains a number of points less than or equal to a predefined threshold. The result is a list of balanced regions represented as bounding boxes.
   - **Benefits of region-based partitioning**
     - **Preserving local spatial relationships**: Each region retains a localized subset of the sample, preserving the spatial context for accurate segmentation.
     - **Enabling efficient parallel processing**: By splitting the data into balanced regions, the pipeline allows for independent processing of each region.

##### C.1.6 Tile creation within each region

For further speed up and following (mini-batching) best practices, once balanced regions are generated, they undergo further processing to create smaller spatial units known as tiles. The main steps are the following:

1. **Data initialization and loading for each region**: For each region, an in-memory dataset is initialized using the spatial boundaries of that region. This dataset is responsible for loading and filtering spatial data to ensure relevant information is retained for further processing.

- **Transcripts loading and filtering**: Transcripts from the `transcripts.parquet` file are loaded based on the region's spatial extents. Transcripts are further refined through quality filtering and removal of unwanted features, such as control probes, according to predefined substrings. Only high-quality transcripts within the specified region are kept for further analysis.
- **Boundaries loading and filtering**: Boundaries either from the `nucleus_boundaries.parquet` or the `boundaries.parquet` file are loaded and filtered to retain boundaries that intersect with or lie within the region's extents. This ensures that spatial relationships between transcripts and boundaries are preserved properly.

2. **Generating tiles within each region**: Each balanced region is subdivided into smaller rectangular tiles using user-specified dimensions for width and height.

    - **Rectangular tile generation**: The region is divided into a grid of rectangular tiles, each represented as a polygon.

3. **Random assignment of tiles to data subsets**: Each generated tile is randomly assigned to one of three data subsets: training, validation, or test. This assignment ensures the unbiased distribution of data for model training and evaluation.

##### C.1.7 Graph construction from tiles

Once the spatial tiles are generated, each tile undergoes conversion into a graph representation using *PyTorch Geometric* (*PyG*) data objects. This process models the interactions between transcripts and boundaries in a graph structure that captures both spatial and functional relationships.

1. **Tile initialization and data extraction**

    - **Initialization**: Each tile is initialized with its spatial extents, and relevant data (transcripts and boundaries) is extracted from the corresponding region's in-memory dataset.
    - **Filtering**: Transcripts and boundaries within the tile are filtered to retain only those within or near the tile's spatial extents, ensuring that only relevant spatial entities are included.

2. **Node Construction**

    - **Transcript Nodes**:
        - `data["tx"].id` : Unique identifier for each transcript.
        - `data["tx"].pos` : Spatial coordinates of the transcript within the tile.
        - `data["tx"].x` : Feature vector representing the transcript, either as a gene-cell-type abundance embedding (if scRNA-seq data is available) or as a one-hot encoding of its feature label.
    - **Boundary Nodes**:
        - `data["bd"].id` : Unique identifier for each boundary.
        - `data["bd"].pos` : Spatial centroid coordinates of the boundary.
        - `data["bd"].x` : Geometric properties, including:
            * **Area**: Surface area of the boundary polygon.
            * **Convexity**: Ratio of the convex hull area to the polygon's area.
            * **Elongation**: Ratio of the minimum bounding rectangle's area to the polygon's envelope area.
            * **Circularity**: Ratio of the polygon's area to the square of its minimum bounding radius.

3. **Edge construction**

    - **Transcript-neighbors-transcript edges**:
        - Represent spatial proximity between transcripts within the tile.
        - Computed using a KD-Tree for nearest-neighbor queries, based on user-specified parameters: number of nearest neighbors ( `k_tx` ) and maximum distance ( `dist_tx` ).
        - `data["tx, "neighbors", "tx"].edge_index` : Tensor storing edge indices for these connections.
    - **Transcript-belongs-boundary edges**:
        - Represents spatial containment where a transcript overlaps with a boundary, indicating that the transcript is expressed within that boundary.
        - `data["tx", "belongs", "bd"].edge_index` : Tensor storing edge indices for these relationships.
    - **Negative edge sampling and labeling**:

- Random negative edges are added using the `RandomLinkSplit` transformation. This creates a balanced set of positive and negative edges, crucial for training the link prediction model. Positive edges denote true relationships, while negative edges represent artificially generated non-relationships.
- Edge labels distinguish positive (true) edges from negative (sampled) edges, supporting a binary classification task where the goal is to predict whether a specific transcript is expressed within a specific boundary.

#### C.2 Model training

The training process for the segger model focuses on identifying which transcripts belong to a specific nucleus or cell type boundary. This connection reflects the biological relationship of transcripts being expressed by specific cells. By framing the task as link prediction, the model learns to predict transcript–cell associations using graph representations of spatial regions.

Training tiles are used to optimize the model, while validation tiles monitor performance during training and determine the best model. Test tiles are reserved for evaluating the model's final performance. The *LitSegger* module leverages PyTorch Lightning to manage data loading, training, and validation, enabling the use of multiple GPUs to significantly accelerate the process. Using binary cross-entropy loss, the model refines its predictions, which are evaluated with the metrics AUROC and F1 Score.

##### C.2.1 Input data

The input data for model training consists of spatial tiles, represented as *PyTorch Geometric* (*PyG*) data objects. These tiles are split into three subsets, each stored in a separate directory:

- ***Training set***: Contains tiles used for model training.
- ***Validation set***: Contains tiles used for monitoring the model's performance during training.
- ***Test set***: Contains tiles reserved for evaluating model performance after training is complete.

The tiles are loaded as `STPyGDataset` objects, which is a subclass of *PyG* `InMemoryDataset`. The `InMemoryDataset` serves as a collection of *PyG* `Data` objects, where each *PyG* `Data` object represents a graph corresponding to a spatial tile. These graphs contain nodes, edges, and their associated features, as described in the Data preprocessing section.

##### C.2.2 Required parameters

Several parameters guide the model training workflow, either specified through the CLI or a YAML configuration file:

- `data_dir` : Directory containing the processed segger dataset (train, validation, and test tiles).
- `models_dir` : Directory to save the trained model and training logs.
- `sample_tag` : Sample tag for the dataset being used.
- `num_tx_tokens` : Number of transcript tokens used for representing transcript features.
- `init_emb` : Size of the embedding layer used for transcript features.
- `num_mid_layers` : Number of middle layers in the model architecture.
- `hidden_channels` : Number of hidden channels in middle layers.
- `heads` : Number of attention heads used in graph attention layers.
- `out_channels` : Number of output channels in the last layer.
- `batch_size` : Batch size for training.
- `num_workers` : Number of workers used for data loading (parallelization).
- `accelerator` : Device type for training (e.g., "cuda" for GPUs or "cpu").
- `max_epochs` : Maximum number of epochs for training.
- `save_best_model` : Whether to save the best model.

- `learning_rate` : Learning rate for training.
- `pretrained_model_dir (optional)` : Directory containing the pretrained model to use (if any).
- `pretrained_model_version (optional)` : Version of the pretrained model.
- `devices` : Number of devices (e.g., GPUs) to use during training.
- `strategy` : Training strategy to be used by the Lightning Trainer.
- `precision` : Determines the numerical precision used during training (e.g., "16-mixed").

##### C.2.3 Data loading using segger data module

`SeggerDataModule` , a subclass of `LightningDataModule` from *PyTorch Lightning*, is used to facilitate efficient data handling for model training, validation, and testing. The data module loads the processed data into three distinct `STPyGDataset` objects, corresponding to the training, validation, and test sets. Data loaders are then defined for each dataset, providing mini-batches of data during training. The size of each mini-batch is defined by the user-specified `batch_size` parameter, and data loading can be parallelized using the `num_workers` parameter.

##### C.2.4 Model training with LitSegger

The `LitSegger` module orchestrates the training and validation of the `segger` model using PyTorch Lightning. The `segger` model itself applies graph attention layers to process heterogeneous graph data, making it suitable for predicting interactions between transcripts and boundaries in spatial transcriptomics data.

1. **Embedding and input handling**
   - **Transcript node embedding**
     - **With gene-cell-type abundance embeddings**: If gene-cell-type abundance embeddings are available, they are used directly as input features for transcript nodes. These features are processed through a linear transformation layer.
     - **Without gene-cell-Type abundance embeddings**: If no such features are available, transcript nodes are represented as tokens and passed through an embedding layer that transforms them into dense feature vectors. The embedding size is defined by the `init_emb` parameter, and the total number of possible tokens is specified by `num_tx_tokens` .
   - **Boundary node transformation**: Features for boundary nodes are processed through a linear transformation layer to match the embedding size of transcript nodes, ensuring a consistent feature space for both node types.
2. **Graph attention mechanism**
   - The `segger` model applies graph attention layers using the `GATv2Conv` operation from *PyTorch Geometric*. This mechanism allows the model to learn weighted relationships between nodes, making it capable of capturing spatial and functional interactions.
   - **Layer structure**
     - **Initial layer**: Processes input features and produces hidden node representations.
     - **Middle layers**: Consist of a configurable number of attention layers ( `num_mid_layers` ) to deepen the model's understanding of complex relationships.
     - **Final layer**: Generates output embeddings for nodes, which are used for the subsequent link prediction task.
   - **Attention heads**: The number of attention heads is specified by the `heads` parameter, allowing multiple perspectives on node connectivity during message passing.
3. **Forward pass and node embedding computation**
   - The forward pass takes node features and edge indices as inputs, propagating them through the layers of the `segger` model to produce node embeddings. These embeddings encapsulate spatial and functional relationships among transcripts and boundaries.
   - After processing, the node embeddings for transcripts and boundaries can be used to predict whether a specific transcript belongs to a specific boundary, forming the basis for the model's link prediction task.

4. **Link prediction and edge scoring**

   - The model's primary task is link prediction, where the goal is to determine if a given transcript is expressed within a particular boundary. This is achieved by computing interactions between transcript and boundary embeddings.
   - **Edge label prediction**: For each edge of type *transcript-belongs-boundary*, the model predicts a score that reflects the probability of a true connection.
   - **Binary classification task**: This prediction is treated as a binary classification problem, with a positive label indicating a true relationship and a negative label indicating no relationship.

5. ***PyTorch Lightning Trainer***

   - The *PyTorch Lightning Trainer* is used to manage the training process. It is configured with the user-specified parameters `accelerator` , `strategy` , `precision` , `devices` , and `max_epochs` .
   - By invoking the `fit()` method, the trainer automatically handles the execution of training and validation steps, including data loading (using `SeggerDataModule` ), forward and backward passes, and optimization.

6. **Training and optimization**

   - The model is trained using binary cross-entropy loss with logits, ensuring that it learns to accurately predict link labels based on the input data.
   - Optimization is handled by the *Adam* optimizer, configured within the `LitSegger` module, with the learning rate set to $1e - 3$ by default.

7. **Metrics and validation**

   - During validation, the model's performance is evaluated using AUROC (Area Under the Receiver Operating Characteristic Curve) and the F1 Score, providing insights into its predictive capability.
   - Validation metrics are logged throughout training, allowing for monitoring of model improvements and potential overfitting.

#### C.3 Segmentation

Segmentation assigns transcripts to boundaries by leveraging the trained `LitSegger` model for link prediction. Through batch-wise processing, transcript-to-boundary similarity scores are calculated, and unassigned transcripts are handled using connected components analysis if enabled. Results are saved in a structured format such as AnnData for downstream analysis. The workflow is designed to be scalable and efficient, with GPU acceleration for computationally intensive tasks and Dask-based parallelized operations for memory-efficient saving and processing of large datasets.

##### C.3.1 Input data

Segmentation relies on the following input data:

- `segger_data_dir` : The directory containing the processed segger dataset, including tiles represented as *PyTorch Geometric* data objects. These tiles retain the graph structure described during data preprocessing and are essential for batch-wise segmentation.
- `models_dir` : The directory containing the model checkpoint saved during training, with the learned weights of the `LitSegger` module.

##### C.3.2 Required parameters

Several parameters guide the segmentation workflow, either specified through the CLI or a YAML configuration file:

- `segger_data_dir` : Path to the processed segger dataset directory.
- `models_dir` : Directory containing the trained model checkpoint.
- `model_version` : Version of the trained model to load.
- `transcripts_file` : Path to the `transcripts.parquet` file.

- `benchmarks_dir` : Directory to save the segmentation results.
- `save_tag` : Tag to use for saving segmentation results.
- `file_format` : Output file format for segmentation results (e.g., *anndata*).
- `cell_id_col` : Column name for storing cell IDs in the output.
- `min_transcripts` : Minimum number of transcripts required for a cell to be included.
- `batch_size` : Batch size for segmentation.
- `num_workers` : Number of workers used for data loading (parallelization).
- `Knn_method` : Method for computing nearest neighbors (e.g., *kd_tree*).
- `score_cut` : The threshold for assigning transcripts to cells based on similarity scores.
- `use_cc` : Whether to use connected components analysis for unassigned transcripts.
- `k_tx` : Number of nearest neighbors for transcript–transcript edges.
- `dist_tx` : Maximum distance for transcript–transcript edges.
- `k_bd` : Number of nearest neighbors for transcript-boundary edges.
- `dist_bd` : Maximum distance for transcript-boundary edges.

##### C.3.3 Initialization

The segmentation process begins with the following initialization steps:

1. **Loading data loaders**: The `SeggerDataModule` , previously used during model training, is reinitialized to load the training, validation, and test dataloaders. These dataloaders enable batch-wise processing of tiles during segmentation.
2. **Loading the model**: The `LitSegger` module is loaded from the specified checkpoint directory. The version of the model to be used is determined by the `model_version` parameter.
3. **Setting up output directories**: A dedicated directory is created to store the segmentation results, organized by the provided `save_tag` . The directory structure includes additional metadata, such as threshold and receptive field parameters, to document the settings used for segmentation.

##### C.3.4 Batch-wise prediction

Once initialized, segmentation proceeds by processing the tiles in batches through the `predict_batch` function, which handles the computations for assigning transcripts to boundaries. This batch-wise approach ensures scalability and efficient memory usage while leveraging GPU acceleration and Dask for parallelized operations. The segmentation steps in `predict_batch` are as follows:

**1. Preparing the batch for GPU processing** Each batch is transferred to GPU for faster computation using CuPy and PyTorch. The model's weights and input data are also loaded onto the GPU to perform similarity score calculations.

**2. Computing transcript-to-boundary similarity scores** The `get_similarity_scores` function computes pairwise similarity scores between transcripts and boundaries. These scores represent the probability of a transcript being expressed within a specific boundary.

**3. Assigning transcripts to boundaries** For each transcript, the boundary with the highest similarity score is identified, provided the score exceeds the user-specified threshold ( `score_cut` ). Transcripts with scores above this threshold are assigned the corresponding cell ID, while those below the threshold are marked as unassigned. The assignments, including transcript IDs, scores, and assigned cell IDs, are stored for further processing.

**4. Handling unassigned transcripts** If enabled ( `use_cc=True` ), unassigned transcripts are grouped using connected components analysis. A transcript–transcript similarity graph is constructed, with edges representing proximity and strong similarity scores. Connected components in this graph are identified to assign cell IDs to unassigned transcripts.

**5. Incremental saving of results with Dask** The transcript assignments are saved incrementally as parquet files for scalability and memory efficiency:

- Transcripts and their computed assignments (e.g., transcript IDs, similarity scores, and assigned cell IDs) are saved to `transcripts.parquet` .
- If connected components analysis is enabled, the transcript–transcript edges used in the grouping process are saved to `edge_index.parquet` .

This incremental saving ensures compatibility with large datasets by avoiding memory constraints.

**6. GPU Memory Management** After processing each batch, GPU memory is cleared using CuPy's memory pool management to ensure resource availability for subsequent batches. This step is critical for processing large datasets without interruptions.

##### C.3.5 Computing similarity scores

To assign transcripts to boundaries, the segmentation process relies on computing similarity scores between nodes in the graph. These scores can be computed for transcript-boundary pairs to determine which transcripts belong to specific boundary or for transcript–transcript pairs to group unassigned transcripts using connected components. The `get_similarity_scores` function performs this computation, leveraging GPU acceleration and the KD-Tree method for efficient nearest neighbor queries. Node types ( `from_type` and `to_type` ) are passed as parameters to specify the type of similarity calculation. The steps are as follows:

**1. Calculating nearest neighbors** The KD-Tree method is used to find the nearest neighbors between the specified node types. The `get_edge_index` function:

- Constructs a KD-Tree using the coordinates of the `to_type` nodes.
- Queries the tree with coordinates of the `from_type` nodes to find up to `k` nearest neighbors within the user-defined distance threshold ( `dist` ).
- Returns the edges as indices in a tensor, representing the adjacency structure for the similarity computation.

**2. Generating node embeddings** The graph neural network processes the graph data to generate embeddings for the `from_type` and `to_type` nodes.

**3. Computing similarity scores** For each edge in the adjacency structure:

- The embeddings of the connected nodes are multiplied to compute a similarity value that quantifies their relationship (e.g., the likelihood of a transcript being expressed within a boundary).
- The similarity values are normalized using the sigmoid function, ensuring all scores are in the range [0, 1].
- The result is a sparse matrix in COO (coordinate) format containing the similarity scores for all valid pairs of nodes.

##### C.3.6 Post-processing and merging results

After processing all batches, the segmentation results are post-processed to finalize transcript-to-boundary assignments. This includes selecting the most confident assignments for each transcript, merging segmentation results with the original transcript data, and grouping unassigned transcripts using connected components analysis (if enabled). The workflow is as follows:

**1. Loading segmentation results** The intermediate segmentation results, incrementally saved during batch processing into `transcripts.parquet` , are loaded using Dask.

**2. Max score selection** To assign each transcript the most reliable prediction:

- For transcripts predicted to belong to a boundary ( `bound = 1` ), the row with the maximum similarity score is identified for each transcript.
- For transcripts predicted to not belong to any boundary ( `bound = 0` ), the row with the maximum similarity score among unbound predictions is identified.

- These indices are combined, prioritizing `bound = 1` assignments when both bound and unbound predictions exist for the same transcript.

The resulting filtered data ensures that each transcript is associated with its most confident prediction. Dask's parallelized operations efficiently handle this grouping and filtering across large datasets.

**3. Merging results with original transcript data** The filtered segmentation results are merged with the original `transcripts.parquet` data to preserve all metadata associated with the transcripts, even for those not assigned to any boundary. This ensures the completeness of the final output. Dask's parallelized merge operations perform this efficiently.

**4. Handling unassigned transcripts using connected components (if enabled)** For transcripts that remain unassigned ( `segger_cell_id` is missing):

1. **Loading edge data:** The transcript–transcript similarity edges, incrementally saved during batch processing into `edge_index.parquet` , are loaded using Dask.
2. **Creating a similarity graph:**
    - Unique transcript IDs involved in the edges are extracted and mapped to numeric indices.
    - A sparse graph is constructed as a SciPy COO matrix, representing transcript-to-transcript connections where edges encode proximity and similarity.
3. **Connected components analysis:**
    - SciPy's connected components algorithm is used to identify clusters of connected transcripts.
    - Each connected component is assigned a unique label, which is mapped back to the original transcript IDs.
4. **Updating assignments:**
    - For unassigned transcripts, their connected component labels are used as temporary `segger_cell_id` .
    - These updates are merged back into the main transcript DataFrame, ensuring that all transcripts, including previously unassigned ones, have associated cell IDs.

The post-processing step ensures that:

- All transcripts are either assigned to a boundary or grouped into connected components for further analysis.
- The final segmentation results are ready for saving in AnnData format or other specified formats.
- Dask is leveraged throughout to enable parallelized operations, ensuring scalability for large datasets.

##### C.3.7 Saving the results

The final step in the segmentation workflow is saving the processed results, ensuring that the output is structured for further analysis. Depending on the specified options, the results can be saved in multiple formats. This step involves saving transcript assignments, generating optional outputs (e.g., cell masks), and logging the segmentation parameters for reproducibility. The key operations are:

**1. Saving transcript assignments to parquet** If `save_transcripts` is enabled:

- The processed transcript assignments, stored in `transcripts_df_filtered` , are repartitioned into 100 partitions to optimize I/O performance.
- The data is saved as a `segger_transcripts.parquet` file in the specified output directory. PyArrow is used as the underlying engine, using efficient serialization with Snappy compression.
- This file includes comprehensive information about all transcripts, including their IDs, similarity scores, assigned cell IDs, and additional metadata.
- This format is well-suited for integration with downstream data processing pipelines.

**2. Saving results in AnnData format** If `save_anndata` is enabled:

- An AnnData object is created using the filtered transcript assignments ( `transcripts_df_filtered` ). This step involves converting the Dask DataFrame into a Pandas DataFrame to meet AnnData requirements.
- The resulting AnnData file, `segger_adata.h5ad` , is saved in the output directory. This format is widely used in single-cell omics and thus compatible with standard analysis tools in the field.

**3. Generating cell masks (if enabled)** If `save_cell_masks` is enabled:

- The `generate_boundaries` function computes spatial boundaries for cells based on the processed transcript assignments.
- These boundaries are saved as a `segger_cell_boundaries.parquet` file. This format supports visualization and spatial analysis of segmented cells.

**Summary of outputs** The saving step results in:

- `segger_transcripts.parquet` : Contains transcript assignments.
- `segger_adata.h5ad` : Stores results in AnnData format for downstream analysis.
- `segger_cell_boundaries.parquet` : Optional cell boundary data for visual analysis.

The saving process ensures that the segmentation results are accessible, well-documented, and compatible with a wide range of analytical tools.

### D Accelerated Computation for Large Graph Datasets in segger

Our computational framework is designed to handle large-scale graph datasets $G = (V, E)$ using distributed and parallel computing. This section details the mathematical and computational strategies of Dask, PyTorch Lightning, CuPy, and Parquet, along with visual illustrations sourced from the internet.

#### D.1 Dask: Distributed Task Scheduling and Data Parallelism

Dask decomposes computations into a Directed Acyclic Graph (DAG) of tasks. For a graph dataset with $n$ nodes, assume we wish to compute a feature $f(v)$ for each node $v \in V$. The overall computation time can be approximated by

$$T_{\text{total}} \approx \max_{j=1,\dots,m} \sum_{t_i \in P_j} t_i,$$

where $P_j$ denotes the set of tasks assigned to worker $j$ among $m$ workers. Dask's dynamic scheduler minimizes $T_{\text{total}}$ via load-balancing, while its distributed data structures allow for out-of-core processing.

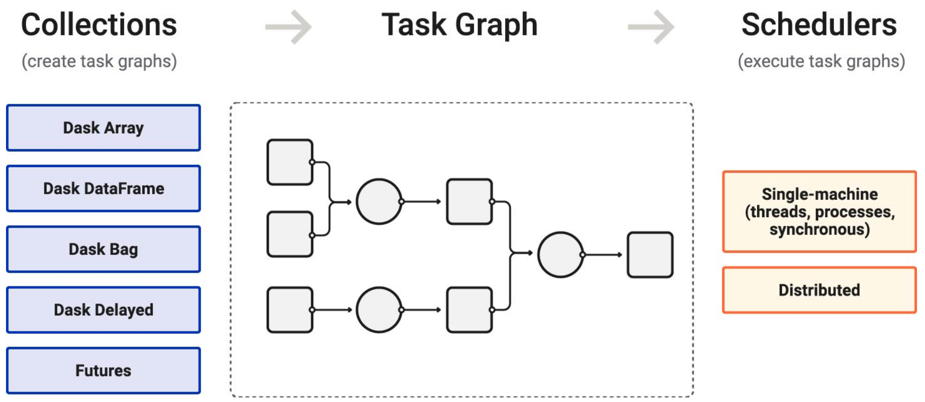

Figure 3: Dask Task Graph Visualization. (Source: https://docs.dask.org/en/stable/10-minutes-to-dask.html)

#### D.2 PyTorch Lightning: Scalable Multi-GPU Training

PyTorch Lightning streamlines deep learning by abstracting boilerplate code and automating distributed training. Consider a graph neural network (GNN) where the loss function $L$ is computed over mini-batches. In a data-parallel setting with $k$ GPUs, each GPU computes a local gradient $\nabla L_i$ so that the aggregated gradient is

$$\nabla L = \frac{1}{k}\sum_{i=1}^{k} \nabla L_i.$$

Thus, the epoch training time is approximately reduced from $C_{\text{GPU}}$ to

$$T_{\text{epoch}} \approx \frac{C_{\text{GPU}}}{k}.$$

Lightning handles gradient synchronization and minimizes communication overhead, making it ideal for high-dimensional graph data.

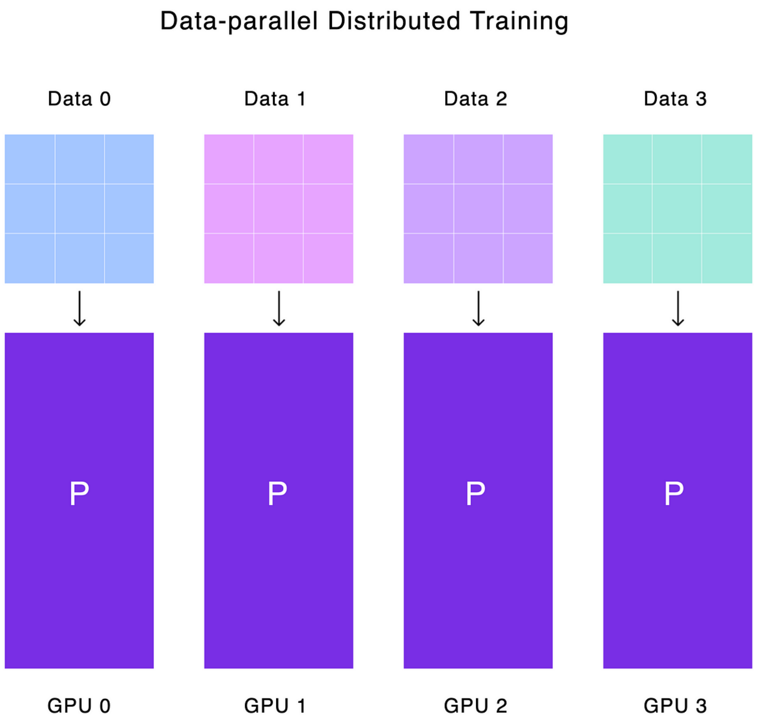

Figure 4: PyTorch Lightning: Distributed Training Architecture. (Source: https://devblog.pytorchlightning.ai/)

#### D.3 CuPy: GPU-Accelerated Numerical Computation

CuPy accelerates array computations on NVIDIA GPUs using CUDA. Many graph algorithms involve linear algebra operations, such as computing the degree matrix $D$ from an adjacency matrix $A$:

$$D_{ii} = \sum_{j=1}^{n} A_{ij}.$$

While this is $O(n)$ per row on a CPU, CuPy parallelizes these operations across thousands of CUDA cores. For example, the matrix multiplication $C = AB$ can be accelerated such that

$$T_{\text{GPU}} \approx \frac{T_{\text{CPU}}}{p},$$

with $p$ representing the effective number of parallel threads.

#### D.4 Parquet: Efficient Columnar Data Storage and I/O Optimization

Apache Parquet is a columnar storage format optimized for large-scale data analytics. For a dataset with $m$ columns, if only a subset $S \subset \{1, \ldots, m\}$ is needed, the I/O cost reduces from $O(m)$ to $O(|S|)$. Formally, the optimized I/O time is

$$I/O_{\text{optimized}} = \sum_{j \in S} \text{size}(C_j),$$

where $\text{size}(C_j)$ is the storage size of column $j$. Parquet's compression and selective read capabilities are vital for fast preprocessing of large graph datasets.

#### D.5 Integration in segger

In segger, these tools are integrated as follows:

- **Dask** orchestrates task distribution and data partitioning across multiple nodes.
- **PyTorch Lightning** automates multi-GPU training, enhancing the scalability of graph neural networks.
- **CuPy** accelerates low-level numerical operations critical for graph analytics.
- **Parquet** optimizes data storage and retrieval, reducing I/O overhead for large datasets.

The combination of these methods minimizes computational overhead and maximizes hardware utilization, making our framework highly efficient for processing large graph datasets.

### E *segger's* step-by-step workflows

**Algorithm 1** Data Preprocessing

**Input:**
- *transcripts_file*: Path to `transcripts.parquet`
- *boundaries_file*: Path to `boundaries.parquet`
- *config_file*: Path to YAML configuration (e.g., `xenium.yaml`)
- *scRNAseq_file* (optional): Path to scRNA-seq data

**Output:**
- *tiles*: Processed spatial tiles saved as *PyTorch Geometric* data objects

**Steps:**

**1. Initialization:**
- Load the configuration file to extract sample type related metadata.
- Initialize directories for training, validation, and test splits.
- Process transcript features:
  - If scRNA-seq data is provided, compute gene-cell-type abundance embeddings and enrich transcript features.
  - Otherwise, represent transcript features using one-hot encodings.

**2. Balanced Region Generation:**
- Compute spatial extents of the sample using boundaries.
- Divide the sample into balanced regions using ND-Tree:
  - Recursively split regions along the largest spatial dimension.
  - Continue until regions meet the size threshold.
- Assign balanced regions for parallelized processing.

**3. Tile Creation:**
- For each balanced region:
  - Load transcripts and boundaries within the region's spatial extents from parquet files.
  - Subdivide the region into smaller tiles of fixed dimensions.
  - For each tile:
    - Filter transcripts and boundaries to include only those within the tile.
    - Assign tiles randomly to training, validation, or test sets.

**4. Graph Construction:**
- For each tile:
  - Create transcript and boundary nodes with their features:
    - Transcripts use gene-cell-type embeddings or one-hot encodings.
    - Boundaries include geometric features like area and elongation.
  - Add edges for transcript proximity using nearest-neighbor queries.
  - Add edges for transcript-boundary overlap (containment).
  - Apply negative edge sampling for transcript-boundary overlap edges.

**5. Save Output:**
- Store graph-structured tiles as *PyTorch Geometric* data objects.
- Save training, validation, and test splits in their designated directories.

**Algorithm 2** Model Training

**Input:** *train_tiles*, *val_tiles*, *test_tiles*
**Output:** *trained_model*: Optimized segger model
**Steps:**
**1. Data Loading:**
Load *train_tiles*, *val_tiles*, and *test_tiles* as PyTorch Geometric datasets.
Create data loaders for mini-batch processing:
    Train loader for model optimization.
    Validation loader for performance monitoring.
    Test loader for final evaluation.
**2. Model Initialization:**
Initialize the segger model with graph attention layers.
Configure transcript and boundary nodes:
    Use gene-cell-type abundance embeddings or one-hot encodings for transcript features.
    Transform boundary features to align with transcript feature dimensions.
**3. Training and Validation:**
Train the model over multiple epochs:
    Perform forward pass through the model to compute transcript-boundary edge scores.
    Calculate binary cross-entropy loss for link prediction.
    Back-propagate gradients and update weights via the Adam optimizer.
    After each epoch, validate the model on *val_tiles* and calculate metrics:
        Area Under the Receiver Operating Characteristic Curve (AUROC)
        F1 Score
**4. Model Selection:**
Identify the model checkpoint with the best validation AUROC and F1 Score.
**5. Save Trained Model and Logs:**
Store the trained model checkpoint for segmentation.
Save training logs for future analysis.

**Algorithm 3** Segmentation

**Input:**

*tiles*: Processed spatial tiles saved as *PyTorch Geometric* data objects

*trained_model*: Optimized segger model

**Output:**

*segmentation_results*: Transcript-to-boundary assignments saved in parquet or AnnData format, optionally including groupings of unassigned transcripts

**Steps:**

**1. Initialization:**

Load tiles for segmentation in batches.
Load the trained model from the checkpoint.
Set up directories for saving segmentation results.

**2. Batch-wise Prediction:**

For each batch of tiles:
- Transfer batch to GPU for efficient processing.
- Compute transcript-to-boundary similarity scores:
  - Generate node embeddings for transcripts and boundaries using the segger model.
  - Calculate similarity scores between transcript and boundary nodes.
- Assign transcripts to boundaries:
  - Assign each transcript to the boundary with the highest similarity score.
  - Mark transcripts with no confident assignments as unassigned.
- Save batch-wise results incrementally.

**3. Handling Unassigned Transcripts (Optional):**

Construct a similarity graph for unassigned transcripts.
Perform connected components analysis to group unassigned transcripts.
Assign temporary cell IDs to grouped transcripts.

**4. Post-Processing:**

Load batch-wise results and merge with the original transcript data.
Select the most confident transcript-boundary assignments:
- Retain the assignment with the highest similarity score for each transcript.

**5. Saving Results:**

Save the final segmentation results:
- Store assignments in parquet or AnnData format.
- Include groupings for unassigned transcripts if applicable.
- Optionally save cell masks for spatial visualization.

[10] Will Hamilton, Zhitao Ying, and Jure Leskovec. Inductive representation learning on large graphs. *Advances in neural information processing systems*, 30, 2017.

[20] T Konstantin Rusch, Michael M Bronstein, and Siddhartha Mishra. A survey on oversmoothing in graph neural networks. *arXiv preprint arXiv:2303.10993*, 2023.

[51] Alexander Kirillov, Eric Mintun, Nikhila Ravi, Hanzi Mao, C. Rolland, and L. Gustafson. Segment anything. *arXiv preprint arXiv:2304.02643*, 2023.
