## Supplementary figures for "Segger: Fast and accurate cell segmentation of imaging-based spatial transcriptomics data"

### Supplementary Tables

**Supplementary Table 1 |** Breakdown of computational costs for segger's workflow on Xenium Breast Cancer dataset, for 3 different embeddings– one-hot (token-based), major (gene-cell-type embedding with 9 cell types), and minor (gene-cell-type embedding with 21 cell types) –across different CPU and GPU configurations, including execution time (seconds) and memory usage (GB). Table provided as ED_table_1.xlsx.

**Supplementary Table 2 |** Coarse cell compartment categories used for Xenium NSCLC dataset annotations, derived from fine-grained cell types in the integrated lung atlas scRNA-seq dataset. Table provided as ED_table_2.csv.

**Supplementary Table 3 |** Consolidated cell type annotations and corresponding marker genes curated from multiple sources, including the Elmentaite et al. colon atlas, a Crohn's disease immune atlas, and 10x annotations of the human colon Xenium panel, with original and re-labeled cell types listed. Table provided as ED_table_3.csv.

**Supplementary Table 4 |** Assignment of 421 genes (FDR < 0.01) to 39 locally co-varying gene modules identified in the segger human colon dataset using Hotspot based on spatial local correlations. Table provided as ED_table_4.csv.

### Extended Data Figures

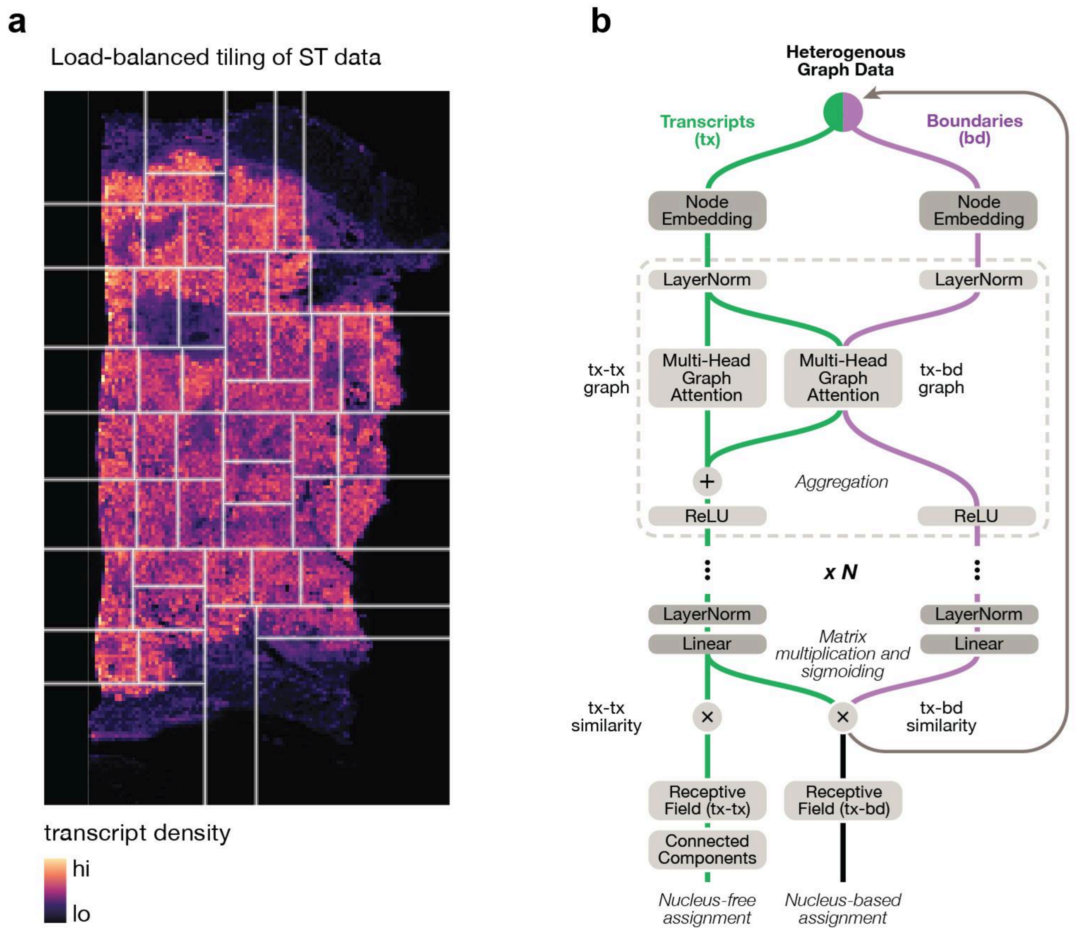


**Extended Data Figure 1.1 | Segger model architecture and adaptive tiling. (a)** Adaptive tiling in segger for scalable model training. The spatial transcriptomics sample is recursively partitioned into load-balanced regions which are processed in parallel. Each region is further subdivided into rectangular tiles of approximately equal size, which are assigned to training, validation, or test subsets for model training. **(b)** Segger graph neural network (GNN) architecture. Segger employs a multi-head graph attention network to propagate features of cell and transcript nodes across k-hop neighborhoods, learning spatial and molecular relationships. Transcripts and boundaries are embedded separately before being aggregated to model intracellular and cell boundary structures. Segmentation is formulated as a binary classification task, predicting transcript-cell associations using a based on the joint latent space and a probit likelihood model. Transcript-Cell assignments are derived within a local field of view, and a predetermined confidence score of association. Transcripts that are not assigned to a nucleus are further grouped into fragments based on transcript-transcript KNN connectivity.

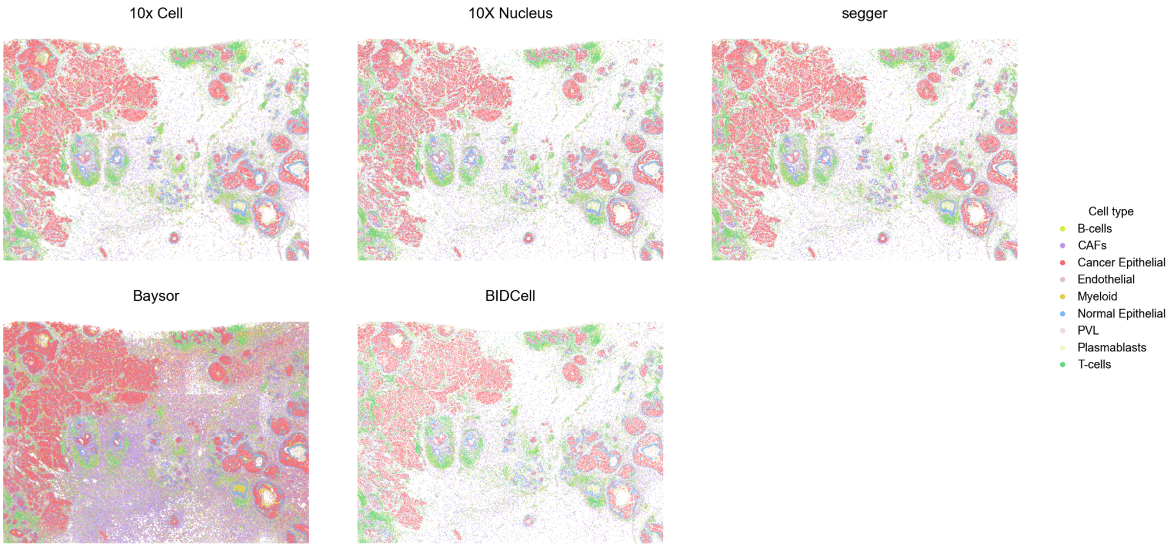


**Extended Data Figure 2.1 | Whole-section cell type annotations for breast cancer Xenium dataset.** Whole-section segmentation and cell type annotations for segger and alternative methods. Cell-type annotations were generated by label transfer from a scRNA-seq breast cancer atlas.

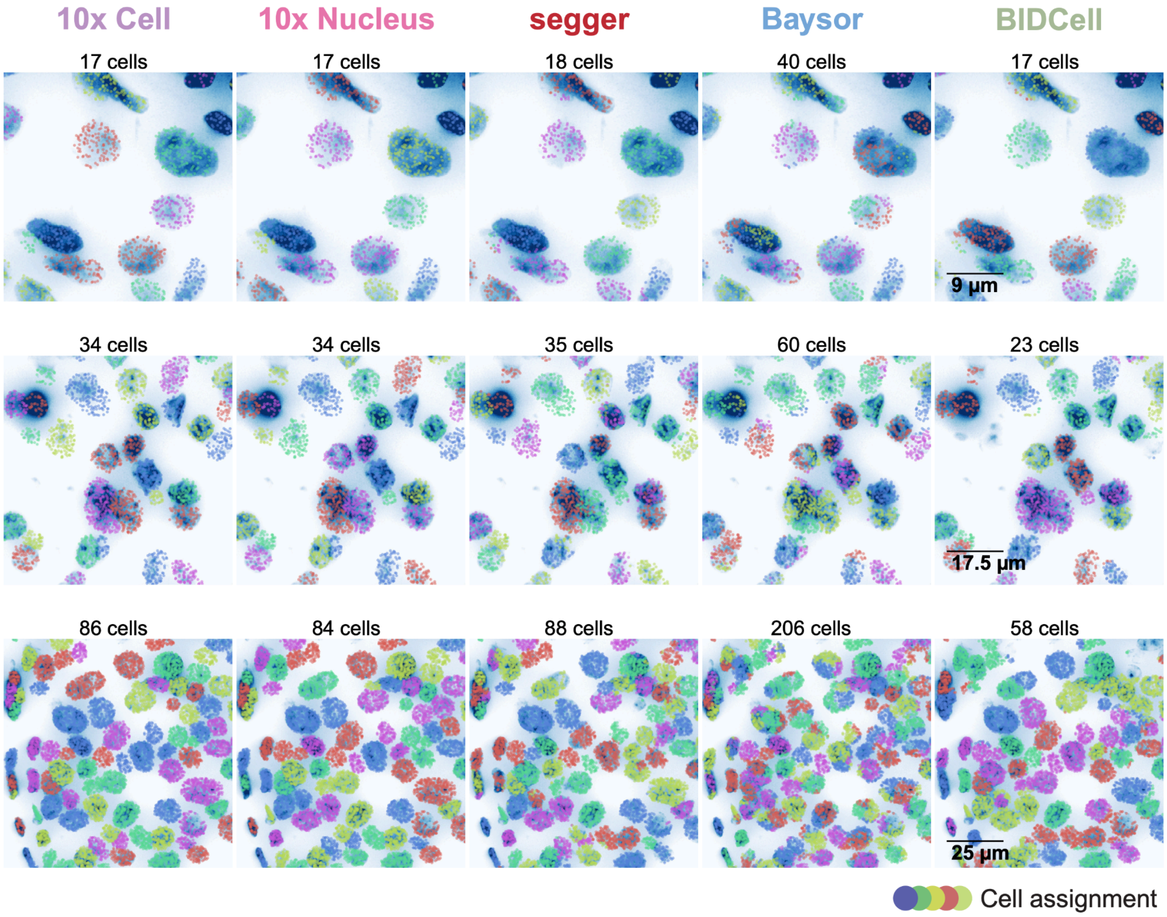


**Extended Data Figure 2.2 | Field-of-view comparisons of cell assignments across segmentation methods.** Representative field of views, showing transcript-to-cell assignments for segmentations generated by segger and alternative methods. Each row corresponds to a different FOV, with transcripts colored by their assigned cell. The number of identified cells is indicated above each panel.

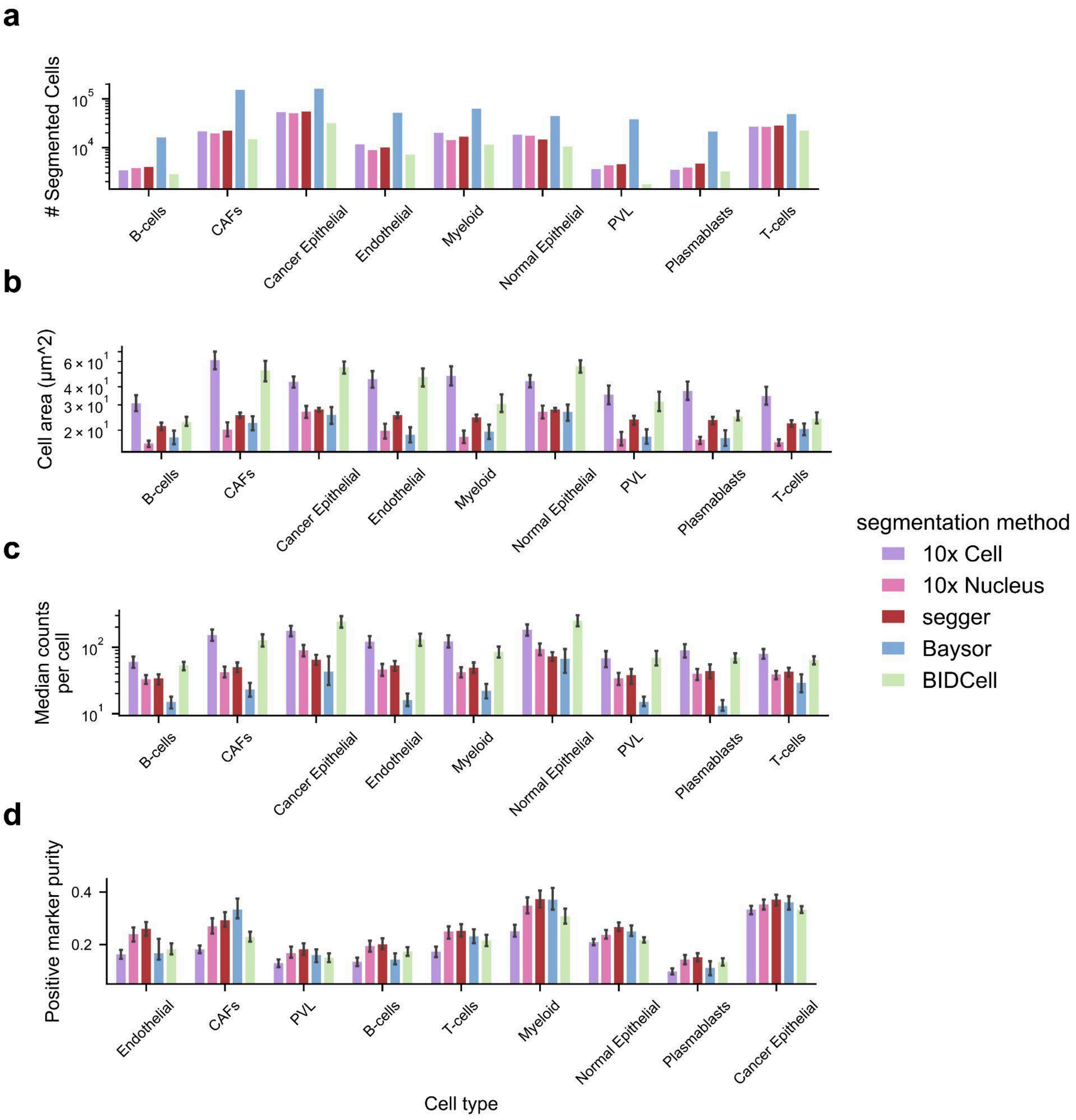


**Extended Data Figure 2.3 | Segmentation metrics stratified by cell type for different segmentation methods in breast cancer Xenium data. (a)** Number of segmented cells per major cell type. (**b)** Median segmented cell area across methods. **(c)** Median transcript counts per cell recovered by each segmentation method. **(d)** Positive marker purity, defined as the proportion of marker-positive transcripts assigned to the correct cell type. Marker genes were derived from a breast cancer scRNA-seq reference dataset.

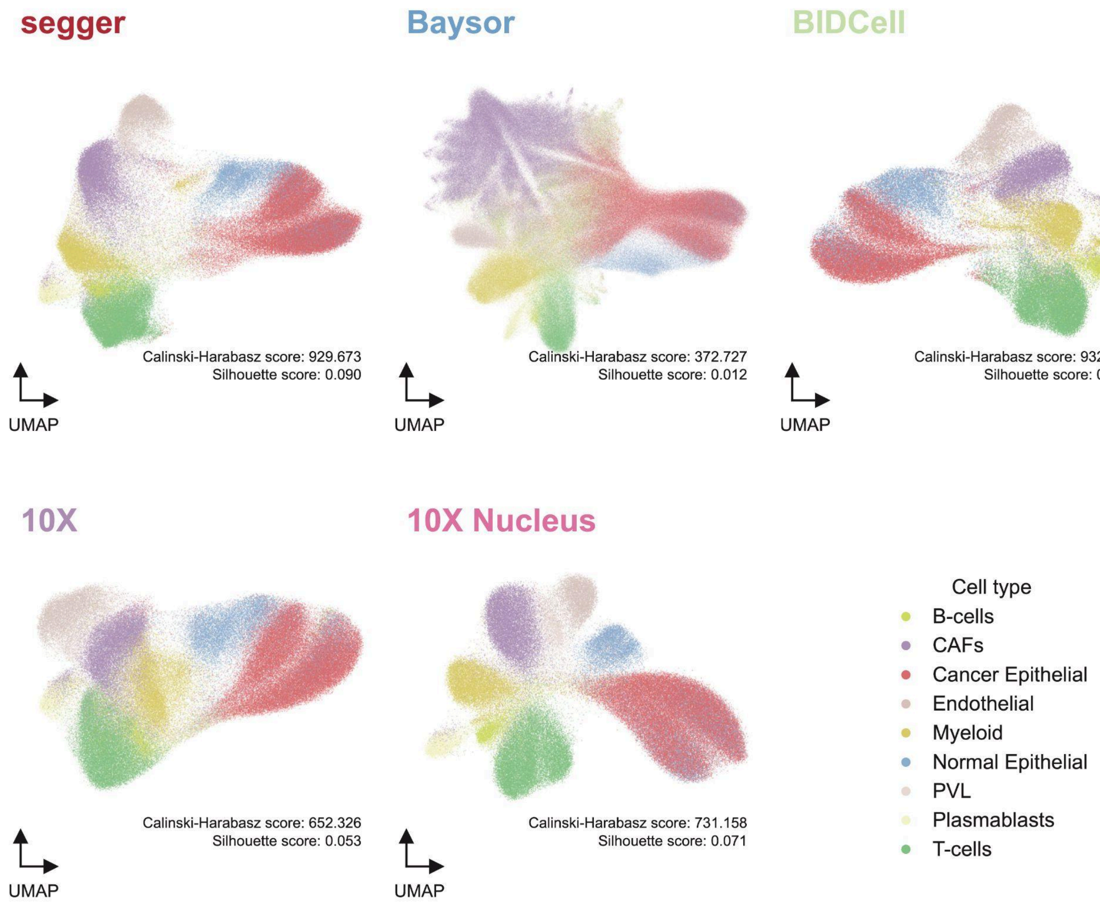


**Extended Data Figure 2.4 | UMAP embeddings of different segmentations for breast cancer Xenium dataset.** UMAP representations of the transcriptome profiles of cells derived using alternative segmentation methods. Points correspond to individual cells, with color corresponding to the cell type assigned using label transfer (**Methods**). Calinski-Harabasz and Silhouette scores are shown for each segmentation method.

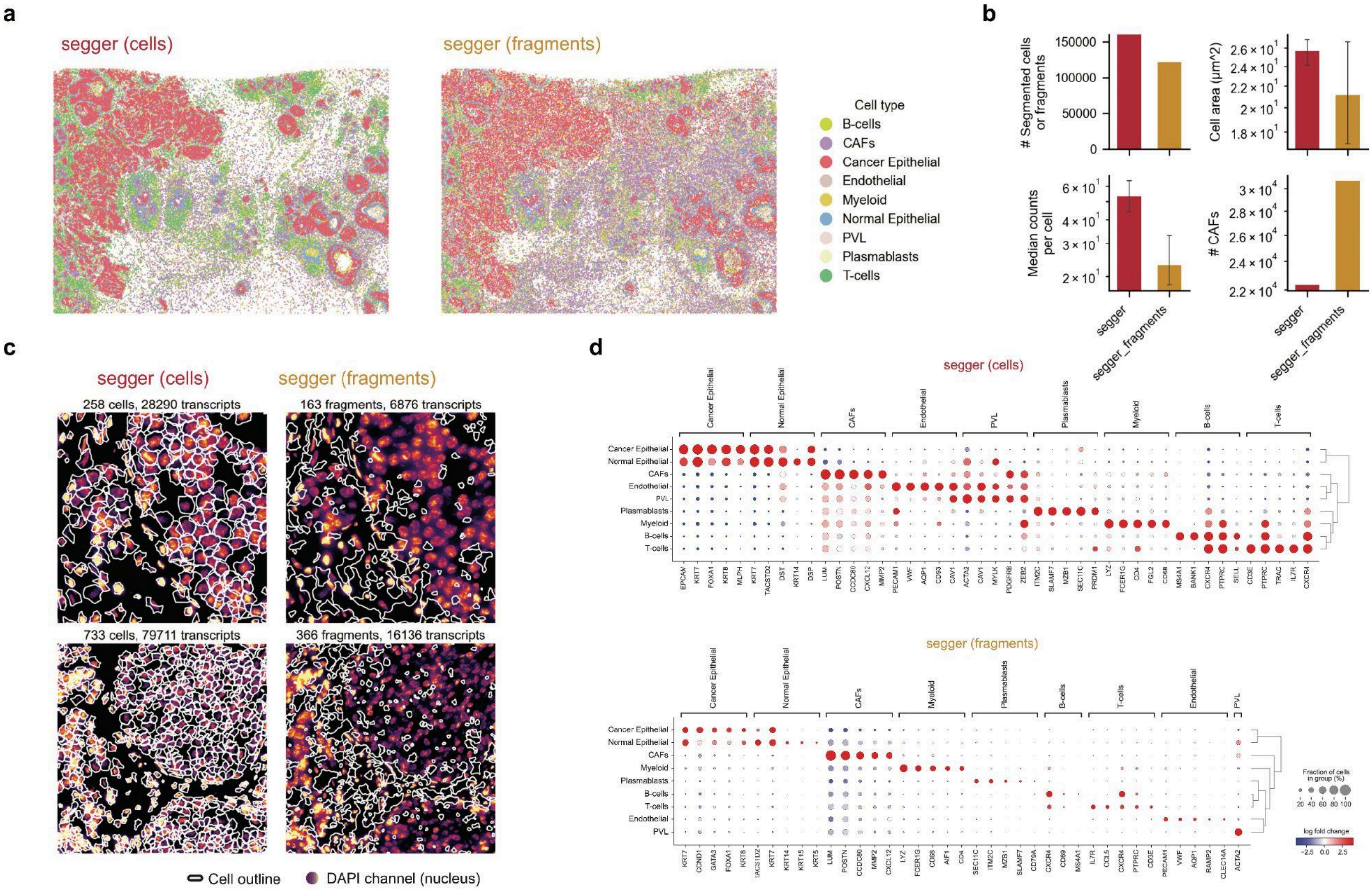


**Extended Data Figure 2.5 | Spatial distribution and marker gene expression of segger cells and fragments. (a)** Whole-section segmentation and cell type annotations for segger-segmented cells (left) and fragments (right) in Xenium breast cancer data. Cell-type annotations were generated by label transfer. (**b)** Selected statistics of segger cells versus fragments: number of segmented componentes, median cell area, median number of counts per cell, and number of CAF-annotated components. Segger fragments are smaller than average cells. Most of the CAF-associated transcripts that are not assigned to any cell by segger, are recovered and grouped into fragments. Error bars represent interquartile range (IQR). (**c)** Segmentation results for two representative fields of views, displaying nucleus staining (DAPI channel) alongside transcripts assigned to cells (left) and fragments (right). The total count of segmented cells and recovered transcripts are displayed above each panel. **(d)** Dot plots illustrating the expression of cell-type-specific marker genes for different cell populations identified in the scRNA-seq reference data, segger-segmented cells, and segger-identified fragments. Each dot represents the expression of a marker gene, with the size of the dot indicating the fraction of cells expressing the gene in the given cell type, and the color indicating the log fold-change relative to other cell types.

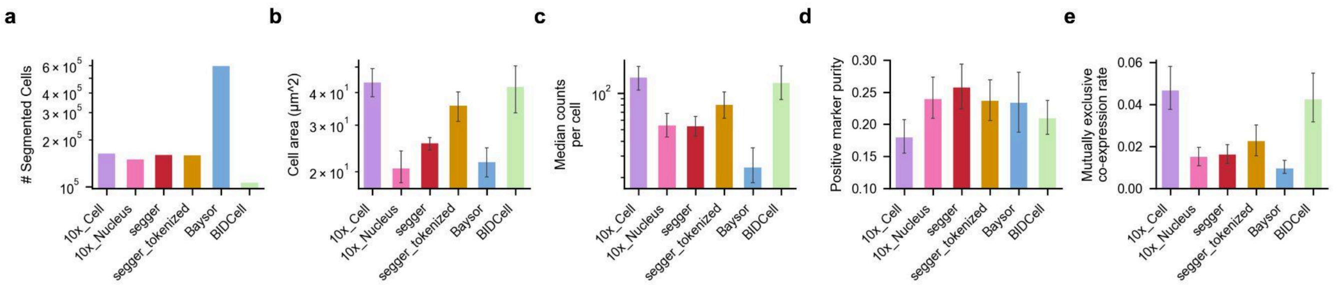


**Extended Data Figure 2.6 | Comparative performance of segmentation methods on the 10x Xenium breast cancer dataset, including segger run either with or without scRNA-seq informed embedding.** Global statistics on segmentation results for alternative methods. Segger corresponds to the full model as considered in Fig. 2, which includes an scRNA-seq guided transcript embedding. Segger_tokenized shows the result when using segger without using such information. **(a)** Total number of cells, with segger and segger-segger_tokenized yielding a comparable number of cells. Cells with less than 5 counts have been discarded. **(b)** Average cell area (μm²), with segger_tokenized yielding slightly bigger cells than segger-embedding. **(c)** Median transcript counts per cell, reflecting the balance between transcript coverage and segmentation. **(f)** Positive marker purity across cells, corresponding to the proportion of correct transcript assignments, as defined from an scRNA-seq atlas. Segger_tokenized achieves overall lower purity than segger, but higher purity values than Baysor, BIDCell, and 10x Cell. **(g)** Mutually exclusive co-expression rate (MECR) for pairs of exclusive transcripts (N=236 gene pairs, <1% codetection rate in parallel scRNA-seq). Shown is, for each pair, the fraction of cells with false-positive co-expression versus cells with exclusive expression. Segger,segger-tokenized, Baysor, and 10x Nucleus achieve comparable MECRs.

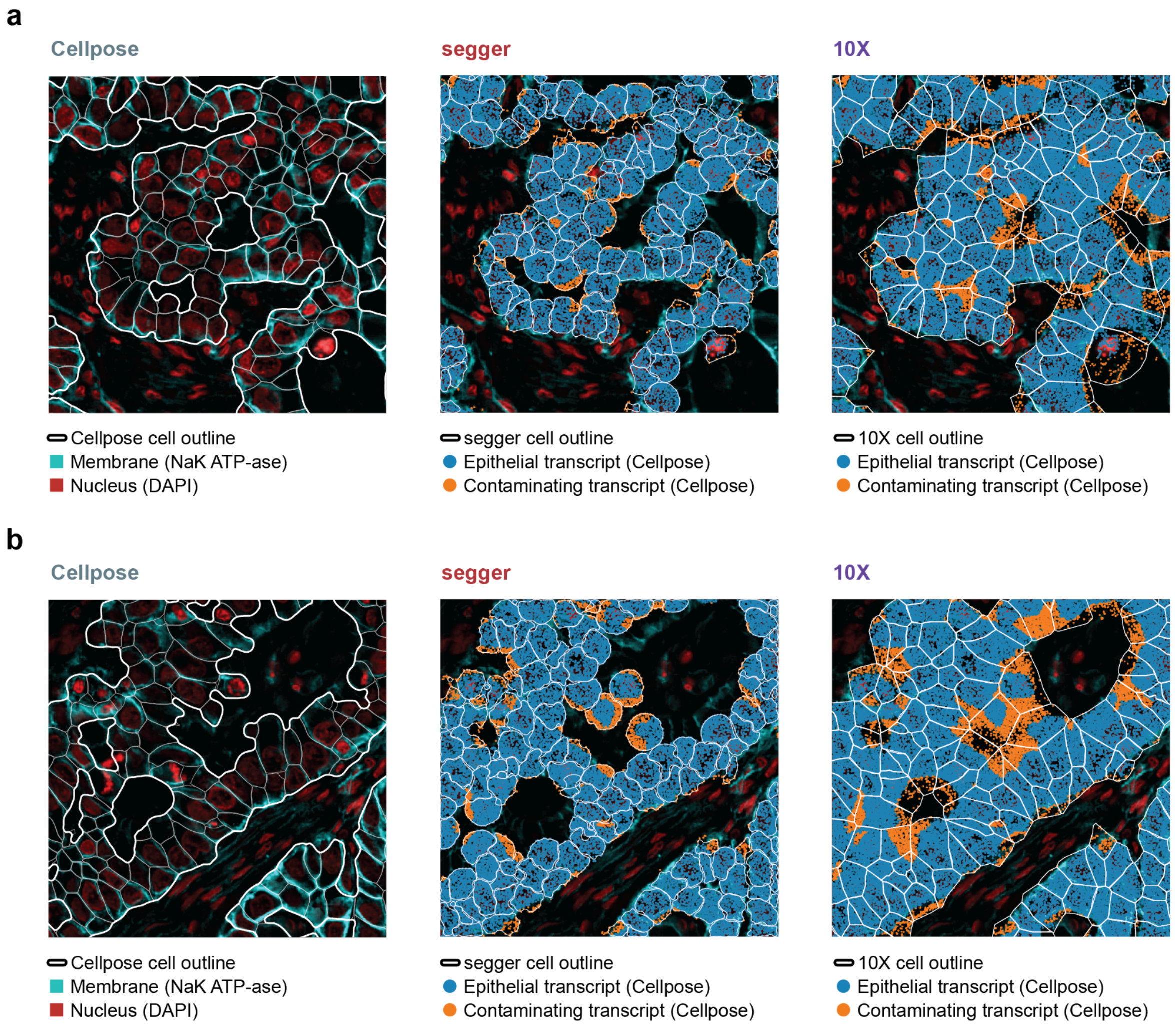


**Extended Data Figure 3.1 | Comparison of 10X and Segger segmentations for additional FOVs.**

**(a, b)** Additional field of views as in Fig. 3, displaying cellpose-segmented cells (left), overlaid with segger (center) or 10x (right) segmented cell boundaries. Cellpose-derived epithelial boundaries serve as a reference, with all segmented epithelial cells outlined in white. Transcripts identified within epithelial cells are colored in blue, while non-epithelial transcripts assigned to epithelial cells (misattributions) are shown in orange. 10X segmentation consistently expands cell boundaries beyond epithelial borders, leading to systematic transcript misassignment. Segger preserves epithelial structures more accurately, reducing contamination.

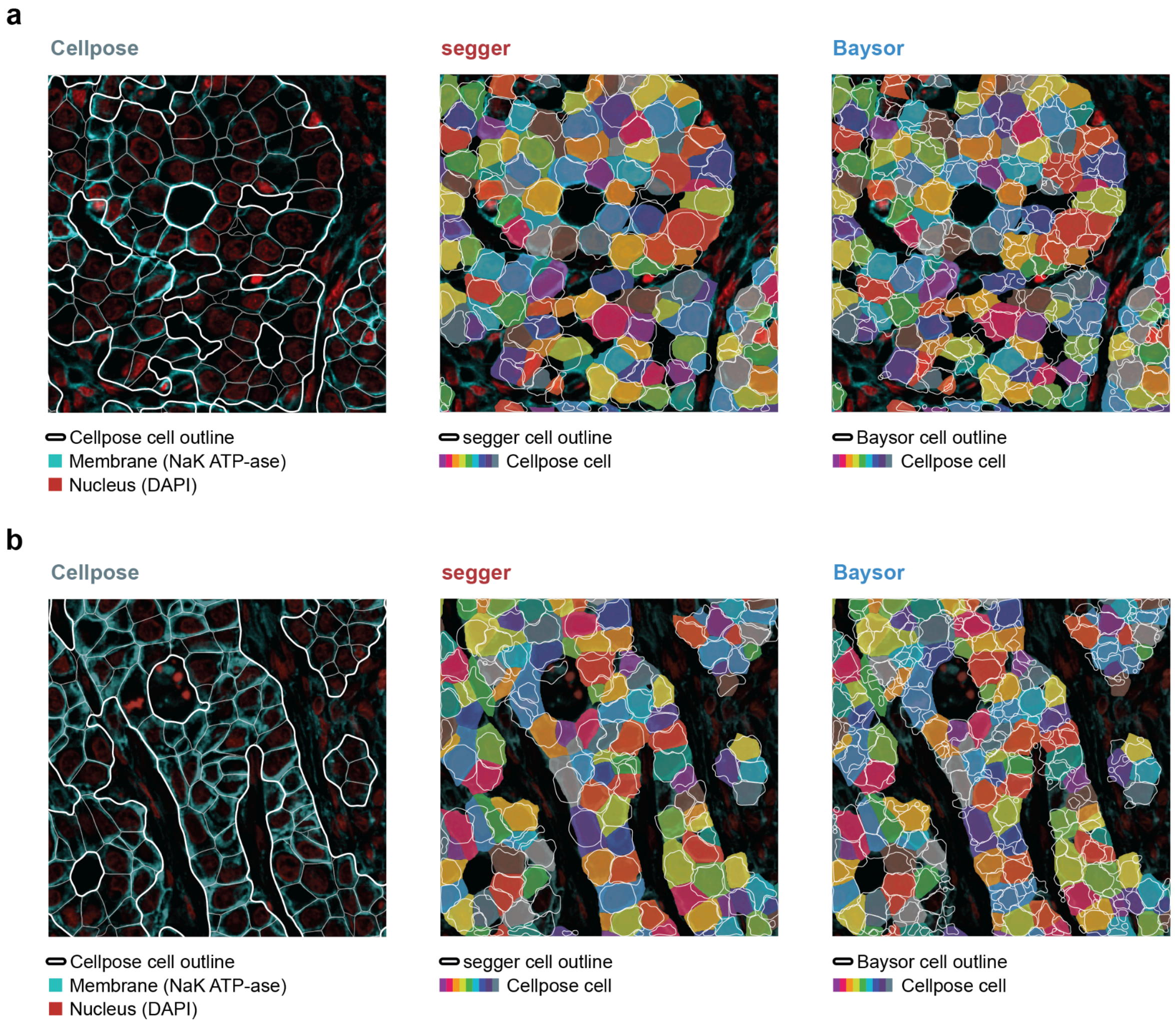


**Extended Data Figure 3.2 | Comparison of 10X and Baysor segmentations for additional FOVs. (a, b)** Field of view (FOV) segmentation comparisons between Cellpose, Segger, and Baysor in two additional lung tissue regions. Cellpose-derived epithelial boundaries serve as a reference, with Cellpose segmented epithelial cells filled with color, and Segger and Basyor cells outlined in white. Baysor consistently over-segments cells resulting in multiple cells per Cellpose cell. Segger preserves cell boundaries more accurately.

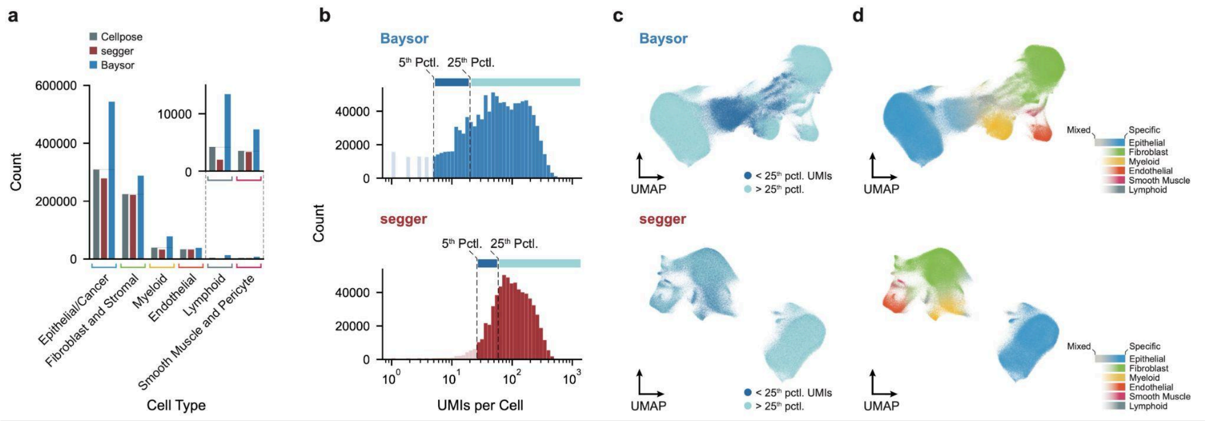


**Extended Data Figure 3.3 | Effect of over-segmentation on transcript density and cell-type separation in UMAP space. (a)** Cell type distribution across segmentation methods. Comparison of the total number of segmented cells per cell type for Cellpose (gray), segger (red), and Baysor (blue). Baysor systematically over-segments, producing an inflated number of cells across all categories, particularly for epithelial and stromal populations. The inset highlights rare cell types, which feature comparable features. **(b)** Transcript density per cell. Distribution of unique molecular identifier (UMI) counts per segmented cell for Baysor (top) and Segger (bottom). Over-segmentation by Baysor results in a large fraction of sparsely populated cells with low transcript counts, as indicated by the large proportion of cells falling below the 25th percentile threshold (light blue). In contrast, segger maintains a more biologically realistic distribution, with the majority of cells containing higher transcript counts. **(c)** UMAP representation of transcript counts. Cells are visualized in UMAP space, colored by transcript count (dark blue: low UMIs; light blue: high UMIs). Baysor produces low-UMI cells spanning high-UMI cells in the embedding, reflecting its fragmentation of true cells into smaller artificial units. Segger preserves higher transcript coverage per cell, resulting in a more structured representation of cell populations. **(d)** UMAP representation of cell type identity. Cells are colored by their assigned cell type (see Methods). Baysor's over-segmentation creates excessive bridging cells between distinct clusters, blurring cell type boundaries and reducing separation between biologically distinct populations. Segger maintains clearer cell type separation, with more well-defined clusters corresponding to epithelial, stromal, myeloid, and endothelial cells. The presence of fewer "mixed" cells in Segger indicates a reduction in artificial cell fragmentation compared to Baysor.

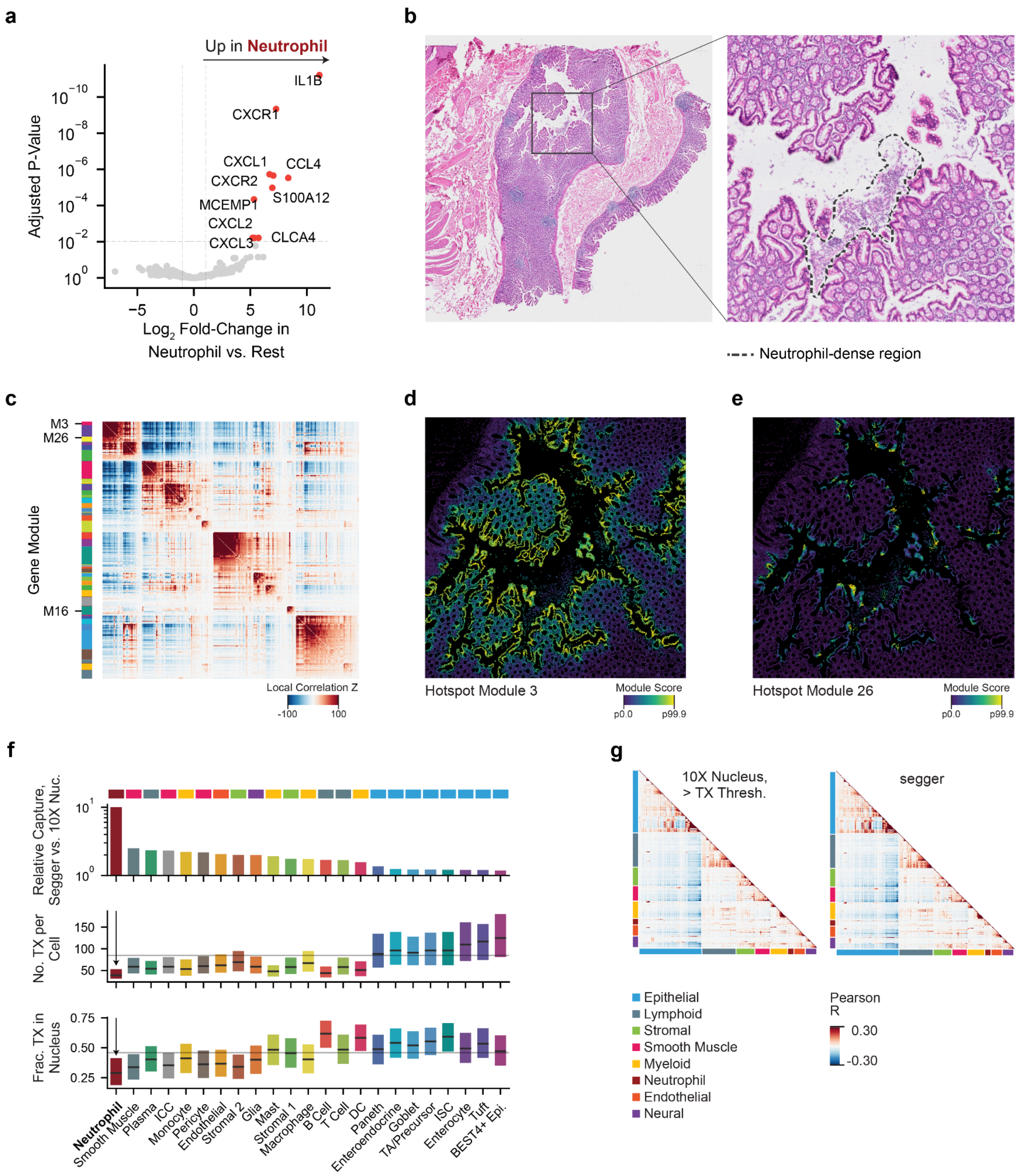


**Extended Data Figure 4.1 | Segger identifies neutrophils in healthy human colon iST data, where nuclear segmentation fails. (a)** Differential gene expression analysis of neutrophils compared to all other cell types in 10x colon Xenium data segmented with segger. Genes with LogFC > 1 and FDR-adjusted $p < 0.01$ are highlighted; Wald test, Benjamini-Hochberg correction. **(b)** Left, hematoxylin and eosin (H&E) stain image collected post-Xenium for an adjacent slide to the 10x healthy colon dataset. Right, a representative neutrophil-dense region is highlighted near the upper-crypt epithelial lining. **(c)** Hotspot modules in all colon cells segmented with segger. The heatmap comprises 421 genes with significant autocorrelation (FDR < 0.01), grouped into 39 gene modules (**Supplementary Table 4** and **Methods**). **(d,e)** Hotspot module scores of cells for module 3 (d) and 26 (e) for the zoomed-in FOV depicted in (b). **f,** Relative cell-type capture rates in segger vs. 10x

Nuclear segmentations (top), transcripts per cell in segger-segmented data (middle), and transcript fraction per cell (bottom), grouped by annotated cell types. In middle and bottom plots, black lines denote median and bars represent interquartile range. Rectangles at top indicate cell compartments, highlighting the lower capture efficiency of non-epithelial cells. **(g)** Gene–gene correlation matrices show that segger, despite having higher sensitivity than 10x Nucleus, provides a similar ability to distinguish cell types with clear correlation structure.
